# Distinct Synaptic Mechanisms Drive *NRXN1* Variant-Mediated Pathogenesis in iPSC-Derived Neuronal Models of Autism and Schizophrenia

**DOI:** 10.1101/2025.07.18.664735

**Authors:** Jay English, Danny McSweeney, Jinghui Geng, Ethan Howell, Fumiko Ribbe, Matthew Hinderhofer, Lydia Proskauer, Rebecca Sebastian, Le Wang, Tal Sharf, Zhiping P. Pang, ChangHui Pak

## Abstract

Copy number deletions in the 2p16.3/*NRXN1* locus confer genome wide risk for autism spectrum disorder (ASD) and schizophrenia (SCZ). Prior work demonstrated that heterozygous *NRXN1* deletions decreases synaptic strength and neurotransmitter release probability in human-iPSC derived cortical glutamatergic induced neurons and this synaptic phenotype is replicated in SCZ patient iPSCs with varying *NRXN1* genomic deletions. What is unknown, however, is whether similar synaptic impairment exists in ASD patients carrying *NRXN1* deletions. Answering this question is important to determine whether all *NRXN1* deletion carriers should be treated similarly or individually, based on their genetic backgrounds and deletion breakpoints. Here, using previously uncharacterized ASD patient iPSC lines, we show that ASD-*NRXN1* deletions impact cortical synaptic function and plasticity in unique ways compared to SCZ-*NRXN1* deletions. Specifically, at a single neuronal level, ASD-*NRXN1* deletions alter basal spontaneous synaptic transmission by selectively enhancing excitatory synaptic signaling with no changes at inhibitory synapses while SCZ-*NRXN1* deletions reduce both excitatory and inhibitory synaptic transmission. At the neuronal network level, there exists enhanced transmission probability and irregular firing patterns in ASD-*NRXN1* deletions. Such changes at the synaptic and network level connectivity patterns influence a critical form of developmental cortical plasticity, synaptic scaling, as ASD-*NRXN1* deletions uniquely fail to upscale their synaptic strength in response to chronic neuronal silencing. Together, these findings highlight the disorder-specific consequences of *NRXN1* deletions on synaptic function and connectivity, offering mechanistic insights with implications for therapeutic targeting and refinement strategies for *NRXN1*-associated synaptopathies.

**Highlights:**

- Novel ASD patient-iPSC-derived E-I culture model to investigate pathogenic *NRXN1* deletions on cortical synaptic function and plasticity
- ASD-*NRXN1* deletions enhance basal synaptic transmission by specifically increasing excitatory neurotransmitter release, opposite of SCZ-*NRXN1* deletions
- Synaptic scaling homeostasis is impaired in ASD-*NRXN1* deletions
- Spike train cross-correlation analysis reveals increased transmission probability and irregular firing patterns in ASD-*NRXN1* deletions

## Introduction

Autism spectrum disorders (ASD) impact ∼3% of the general population in the U.S. and pose a significant burden to caregivers worldwide. ASD arises from hundreds to thousands of genetic variants ranging from single nucleotide polymorphisms (SNPs) in both coding and non-coding regions of the genome, loss-of-function mutations in coding genes, and larger copy number variations (CNVs) that either duplicates or deletes a chromosomal copy. Associating molecular functions onto these ASD risk variants has revealed that certain biological processes might be more vulnerably impacted in autism, thereby giving clues as to how we can approach mechanistic studies in model systems. Notably, synaptic homeostasis, excitatory-inhibitory (E-I) balance and regulation of gene expression are seemingly enriched in the functional categorization of ASD risk genes^1–3^. Therefore, it is not surprising to find multiple genes that encode for synaptic adhesion, scaffolding, and signaling molecules (*NRXN1, NGLN1/3/4X, SHANK2/3, CASK, TSC1/2*) to be discovered as ASD risk genes^1–3^.

*NRXN1* encodes an evolutionarily conserved presynaptic cell adhesion molecule that plays critical roles in synapse specification and maintenance^4^. Neurexin genes (*NRXN1-3*) are extensively alternatively spliced thereby allowing unique protein-protein interaction schemes at the synapse, endowing exquisite functional synaptic diversity throughout the brain^4,5^. Notably, the majority of the heterozygous *NRXN1* mutations that have been identified in patients with ASD and schizophrenia (SCZ) are non-recurrent CNVs that produce heterozygous null mutations in only *NRXN1* due to the large size of this gene, while missense and truncation mutations are less frequent^1,6–15^. Although *NRXN1* gene mutations are rare (∼0.18% of cases across disorders compared to ∼0.02% in controls^1,8,9^), this frequency impacts thousands of patients worldwide. In particular, *NRXN1* deletions show seemingly high odds ratios of 14.4 for SCZ^7^ and 14.9 for ASD^6^. As a result, *NRXN1* mutations represent one of the most frequent single-gene mutations for SCZ and *NRXN1* is currently rated as a SFARI score 1 gene for ASD.

Modeling *NRXN1* deletions in human pluripotent stem cell derived induced cortical excitatory neurons (Ngn2-iNs) revealed that isogenic *NRXN1* heterozygous cKO alleles (exons 18 and 24 deletions) and SCZ patient iPSCs bearing *NRXN1* deletions collectively reduce glutamatergic neurotransmitter release probability and synaptic strength, findings that were replicated across sites in a consortium study^16,17^. Cortical organoids differentiated from the same set of stem cell lines displayed reduction in calcium signaling and network connectivity patterns in *NRXN1* deletion cases^18^. Additionally, in a separate study, Ngn2-iNs bearing *NRXN1* deletions from childhood onset SCZ and Bipolar Disorder patients also showed decreased neuronal signaling^19^. While these studies show agreement in dampened synaptic responses and decreased network function in SCZ patient derived neurons, how the same genetic variant, in our case *NRXN1* deletions, mechanistically act in different disorder contexts is unclear. In other words, can we test and model the pleiotropic nature of *NRXN1* deletions in different disorder contexts? Some clues exist in pointing to increased calcium signaling and neuronal excitability in *NRXN1* deletion carrying ASD patients^20,21^, a phenotype that is in opposition to what has been reported in SCZ patient iPSC studies so far. Moreover, due to the inherent rare nature of the *NRXN1* deletions, there is a general scarcity of patient iPSC lines which limit our overall understanding of the pathophysiological mechanisms underlying this important genetic variant implicated in various neurodevelopmental disorders.

To address these limitations and to gain a detailed mechanistic understanding of how *NRXN1* deletions in ASD background influences synaptic function and connectivity, we partnered with SFARI Stem Cell Initiative to generate a novel set of human iPSC lines from three ASD individuals with specific *NRXN1* mutations (see **Supp. Table 1, Figure 1a,b**). By pairing these three uncharacterized iPSC lines with recently characterized sex-matched healthy control iPSCs, we generated a mixed excitatory (E) and inhibitory (I) induced neuronal cultures at an 80%:20% ratio (E-I iNs), mimicking *in vivo* E-I ratio in the neocortex^22^. This culturing set-up allowed us to study the resulting changes in E-I balance and synaptic transmission in ASD-*NRXN1* deletion cases and how they compare to the established SCZ-*NRXN1* deletions cases^16^.

**Figure 1.**
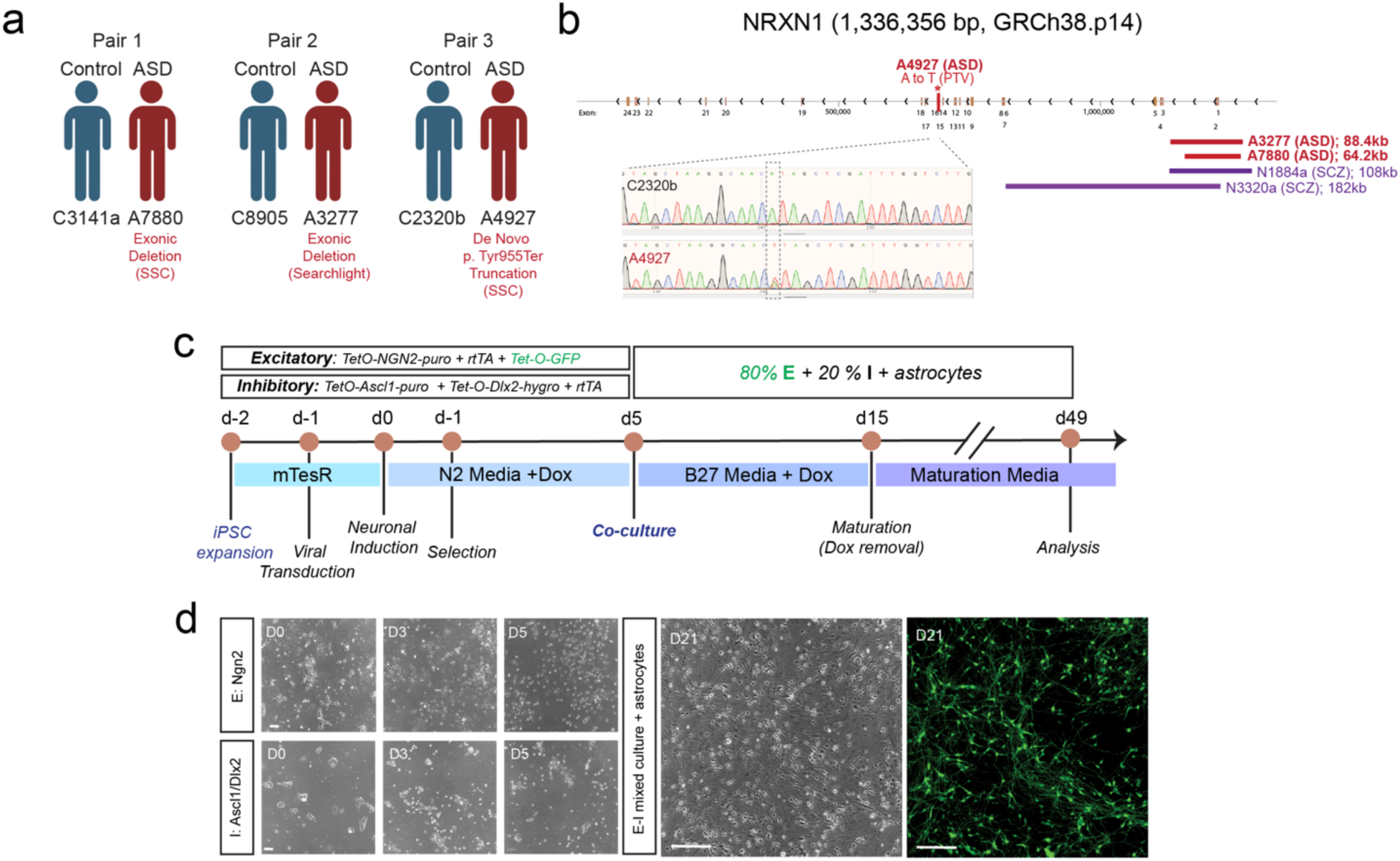
Experiment design for characterizing *NRXN1*^+/-^ ASD patient-derived induced neuronal E-I mixed cultures. **a)** Patient-control paired culturing scheme. ASD patient iPSCs (denoted with an “A” before the identification number) were randomly paired with a sex-matched control neurotypical donor iPSC line (denoted with a “C”). Note all iPSC lines are male. Each pair was differentiated, cultured, and recorded in tandem with each other for each experiment throughout the paper. **b)** Genomic map of the *NRXN1* locus (5’ to 3’ going from right to left) showing the breakpoints of each *NRXN1* deletion/mutation present in individual deletion carrier with ASD (this paper) and SCZ diagnoses (N1884, N3220a; Pak et al., 2021). Individual point mutation for A4927 was confirmed via Sanger sequencing (inlet). See Methods for more details on the corresponding genomic coordinates. **c)** Directed differentiation and culture scheme used for E-I iNs. Human iPSCs were transduced with the indicated lentiviruses for *Ngn2* (top box) and *Ascl1/Dlx2* (bottom box). At 0 dpi (days post-induction), doxycycline was added to the media to turn on expression of transcription factors. At 1 dpi, the indicated antibiotics were added to media for selection (puromycin and/or hygromycin). At 5 dpi, cells were lifted and frozen, and eventually thawed and re-plated in plating media with 80% excitatory, 20% inhibitory cells (on monolayer mouse astrocytes). Cultures were maintained in maturation media (without dox) until the experiments were performed (42-49 dpi). **d)** Representative images of E-I iN cultures throughout the differentiation process (left half) and the maturation process (right half). Excitatory neurons within E-I iNs were labeled with TetO-EGFP during transduction, and so appear green in the mixed E-I iN cultures. Scale bars 100 um on smaller left panels and 200 um on larger right panels.

Using whole cell patch clamp electrophysiology, we find that while intrinsic and active neuronal properties are unaffected overall, ASD-*NRXN1* deletions impair basal synaptic transmission by selectively increasing the frequency of spontaneous glutamate release without changing GABAergic synaptic transmission. Specifically, we observe more frequent fast spontaneous synaptic events, an increased frequency of excitatory miniature synaptic release, and a higher probability of excitatory neurotransmitter release. This phenotype is opposite to SCZ- *NRXN1* deletions, where both glutamatergic and GABAergic synaptic responses are dampened overall, in agreement with previous findings in homogeneous glutamatergic cultures. ASD-*NRXN1* deletions do not exhibit changes in dendritic morphology or synapse numbers, indicating synapse formation is unaltered. In a chronic neuronal silencing paradigm with 48-hr tetrodotoxin (TTX) treatment, ASD-*NRXN1* deletions fail to upscale their synaptic strength, demonstrating a defect in homeostatic synaptic plasticity. These functional changes at the synaptic level translate to an impairment in neuronal network connectivity as evidenced by irregular firing patterns observed in extracellular recordings using high-density microelectrode arrays. Thus, we conclude that *NRXN1* deletions contribute to E-I imbalance by altering synaptic connectivity in distinct and disorder- specific manner.

## Results

### Establishing NRXN1+/- ASD patient-derived E-I iNs

To develop a co-culture system that enables precise control of glutamatergic excitatory (E) and GABAergic inhibitory (I) cortical neuronal densities, we directly differentiated human iPSCs into E-iNs^23^ and I-iNs^24^ in separate experiments and co-cultured the two iN subtypes 5 days post- induction^25^ (**Figure 1c,d**). A similar mixed culturing scheme has previously been successfully applied to synaptic physiology although the ratio of E:I iNs varied slightly (60% E: 40% I)^26^. We chose 80% E: 20% I based on prior studies reporting neuronal densities in the developing neocortex^22^. In addition, we specifically labeled the E-iNs with EGFP to distinguish these cells when performing whole cell patch clamp electrophysiology (**Figure 1d**). In this experimental design, EGFP-labeled E-iNs in the E-I iN co-cultures were patched and analyzed for their membrane properties and synaptic transmission (**Figure 2a**). Hereafter, we refer to these EGFP- labeled excitatory iNs as ‘iNs’ for short unless otherwise noted. Using this culturing scheme, we differentiated three male ASD patient iPSC lines and three male healthy control iPSC lines, whereby an ASD patient line and a healthy control line (passage- and sex-matched) were randomly chosen for pairing. This pairing method ensured that all culturing, differentiations, and electrophysiology experiments were done on the same days between patient and control lines.

**Figure 2.**
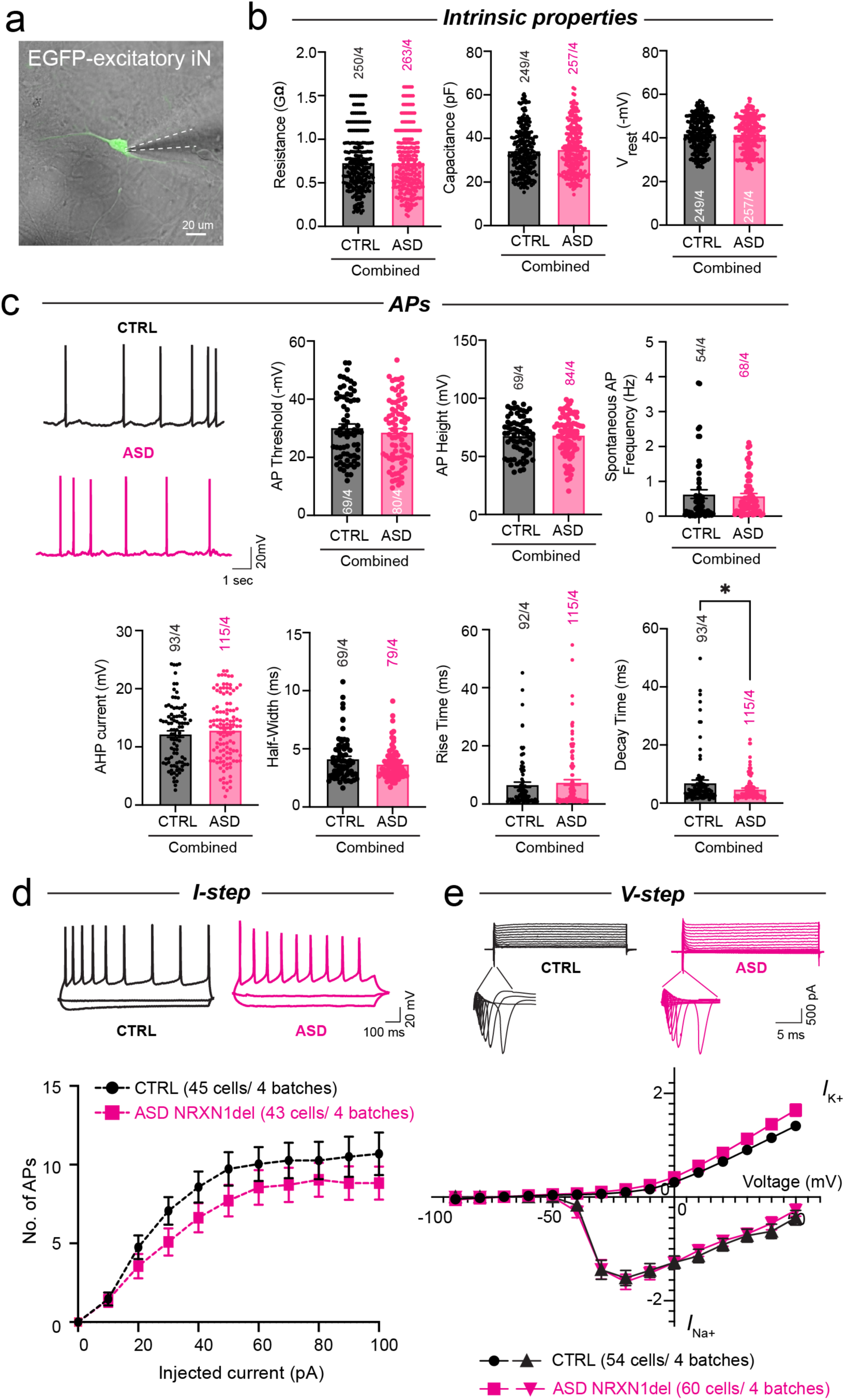
*NRXN1*^+/-^ ASD patient E-I iNs do not exhibit differences in intrinsic properties, action potentials, and sodium/potassium currents compared to controls. **a)** EGFP-positive *Ngn2* excitatory neurons in E-I iN mixed cultures were patched from 3 *NRXN1^+/-^* ASD vs. 3 control iPSC lines and data were pooled and averaged between genotypes. **b)** Summary graphs of measurements of membrane resistance, capacitance, and resting membrane potential of patched excitatory iNs from *NRXN1^+/-^* ASD vs. control E-I iN cultures. No overall change in passive membrane properties was observed. **c)** Summary graphs of spontaneous action potential (AP) properties in *NRXN1^+/-^* ASD vs. control. No differences in AP threshold, amplitude, or frequency were observed. After hyperpolarization potentials (AHP), half-width, and rise time all exhibit no change. A significant decrease in AP decay time was observed in spontaneous AP recordings. **d)** Neuronal excitability is unchanged. Intensity-frequency plot showing the average number of APs as function of step current injections (from −50 pA to +80 pA) in current clamp mode (I-step). **e)** Voltage-dependent Na^+^ (*I*_Na+_) and K^+^ (*I*_K+_) currents are unchanged. Currents were recorded in voltage-clamp mode (V-step). Cells were held at −70 mV while stepwise depolarization was delivered from −90 mV to 50 mV at 10 mV intervals. All summary data are means +/- SEM (numbers in bars or parentheses show number of cells/independent cultures analyzed). Cells were recorded at 42-49 dpi. Statistical analysis was performed with Student’s T-test (*p<0.05).

### NRXN1+/- ASD patient-derived E-I iNs elicit fast spontaneous synaptic transmission without altering intrinsic and active membrane properties

Given that all three ASD patient iPSC lines were previously uncharacterized, we performed a comprehensive set of electrophysiological characterizations. First, we examined their intrinsic properties and generation of spontaneous action potentials (**Figure 2b,c**). At 6-7 weeks post-induction, compared to control iNs, *NRXN1*+/- ASD iNs did not exhibit differences in membrane resistance, capacitance, and resting membrane potentials (**Figure 2b**) or in their ability to generate spontaneous action potentials (APs). Among the seven parameters related to APs quantified, including threshold, height, frequency, half-width, rise time, decay time and after hyperpolarization potentials (AHPs), we only observed a difference in a single parameter - decay time (**Figure 2c**) - in the ASD condition vs. control, potentially indicating faster repolarization speed in these neurons. Furthermore, when we characterized their neuronal excitability by recording the number of APs generated as a function of step current injections, there was no difference between ASD and control conditions (**Figure 2d**). Finally, we examined voltage-dependent sodium and potassium currents with step depolarization protocols and found no difference between the genotypes (**Figure 2e**). All these results suggest that *NRXN1*+/- ASD iNs display comparable passive and active membrane properties and intrinsic neuronal excitability as their control counterparts.

Even though non-synaptic properties are largely intact, changes in synaptic transmission can occur and this phenomenon has been observed in multiple human iN models of ASD risk variants^17,26,27^. To assay synaptic transmission in *NRXN1*+/- ASD iNs, we first measured spontaneous postsynaptic currents (sPSCs) which are mediated by both single vesicular release and network driven synaptic release events including both excitatory and inhibitory synaptic releases. Interestingly, we found that *NRXN1*+/- ASD iNs showed trends toward an increase, albeit no statistical significances, in the average sPSC frequency and variation in the cumulative probability of the inter-event-intervals (IEI) (**Figure 3a,b**). This effect was consistently observed in the pooled summary analysis of all three ASD patient and control lines as well as in the individual analyses by pairs (**Figure S1**). To further investigate deviations in the distribution of sPSCs, we performed additional analysis on sPSC IEIs and noted several differences in the *NRXN1*+/- ASD iNs compared to control iNs (**Figure 3c-f**). First, ASD patient iNs exhibited a significantly smaller range in the total IEI events measured (**Figure 3c**) coupled with an increased frequency of events with shorter IEIs (**Figure 3d**). Second, when we partitioned all IEIs into two different groups based on the histogram distributions (**figure 3d**) – ‘fast’ with IEIs <200 ms intervals and ‘slow’ with IEIs >200 ms intervals – there was a significantly greater percentage of ‘fast’ IEIs with a smaller percentage of ‘slow’ IEIs (**Figure 3e, f**). Interestingly, a similar finding has recently been reported in an *Fmr1* KO mouse model of Fragile X Syndrome, a syndromic form of autism, wherein increased excitatory signaling corresponded with an overall decrease in sEPSC IEIs, particularly exhibiting a dramatic increase in the sub-200 ms range of events^28^. Overall, these findings indicate that ASD patient iNs display more frequent ‘fast’ spontaneous synaptic transmission compared to controls, reflecting enhanced synaptic communication.

**Figure 3.**
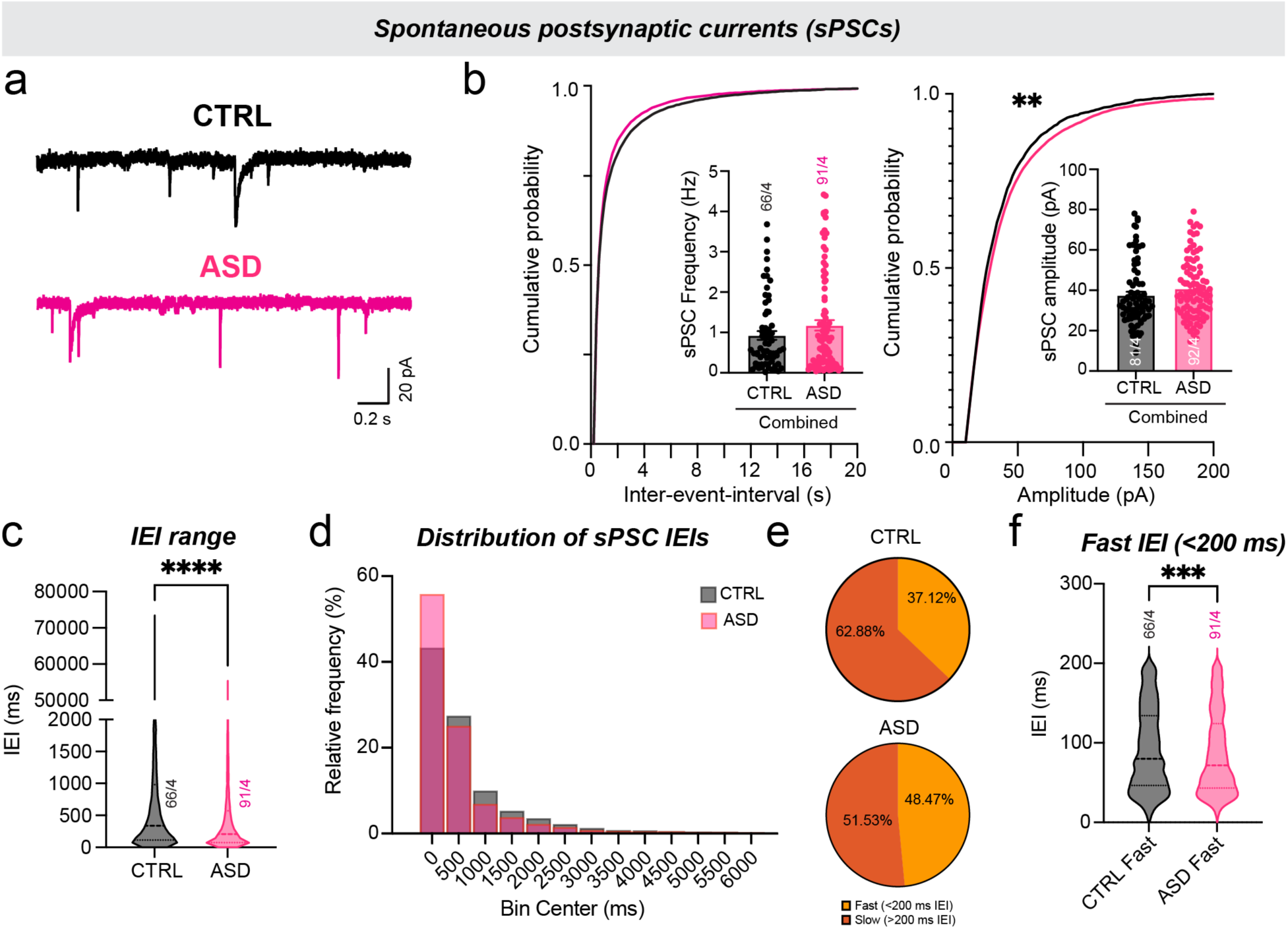
Increased spontaneous synaptic transmission in *NRXN1*^+/-^ ASD patient E-I iNs. **a)** Representative sPSC traces. **b)** Pooled summary graphs of *NRXN1*^+/-^ ASD patient and control recordings of spontaneous postsynaptic current (sPSC) frequency (left) and amplitude (right). Quantifications are shown both as cumulative probability plots of inter-event intervals (IEI) and amplitude as well as average bar graphs (inserts). **c)** Violin plots showing the total range of IEIs among sPSC events with median values (large dots) shown with upper (75%) and lower (25%) quartiles (small dots). **d)** Frequency distribution of sPSC events under each IEI bin (all IEI events below 6000 msec included). **e)** Pie charts showing the proportion of fast (IEI <200 ms) and slow (IEI >200 ms) events out of total sPSC events. **f)** Violin plots showing the median (large dots) IEI (msec) of fast IEI events with upper (75%) and lower (25%) quartiles (small dots). All summary data from panel b are means +/- SEM (numbers inside or above bars show number of cells/independent cultures analyzed). Cells were recorded at 42-49 dpi. Statistical significance for average frequency and amplitude were determined with Student’s T-Test, whereas significance for cumulative probability distributions was determined by two-tailed Kolmogorov-Smirnov test. P-values: * < 0.05, ***<0.001, ****<0.0001.

### Neurite outgrowth and synapse formations are not altered in NRXN1+/- ASD patient-derived E-I iNs

To test whether neuronal morphological complexity is altered in *NRXN1*+/- ASD iNs, we quantified dendritic outgrowth and branching, as well as soma size in the excitatory EGFP-positive iNs from which we recorded. We observed no difference between ASD patient and control iNs (**Figures 4a**, **S2**). This finding recapitulates previous studies in Ngn2-iNs with *NRXN1*+/- SCZ and *NRXN1*+/- cKO alleles^16,17^. In addition to morphological analysis, we performed synaptic density measurements on the same EGFP-labeled dendrites by immunolabeling both excitatory and inhibitory presynaptic puncta, VGLUT1 and VGAT respectively. Similarly, in agreement with previous reports^16,17^, we did not detect any differences in the density and volume of excitatory and inhibitory synapses (**Figure 4b**, **S3**), indicating that synapse formation is normal in *NRXN1*+/- ASD iNs.

**Figure 4.**
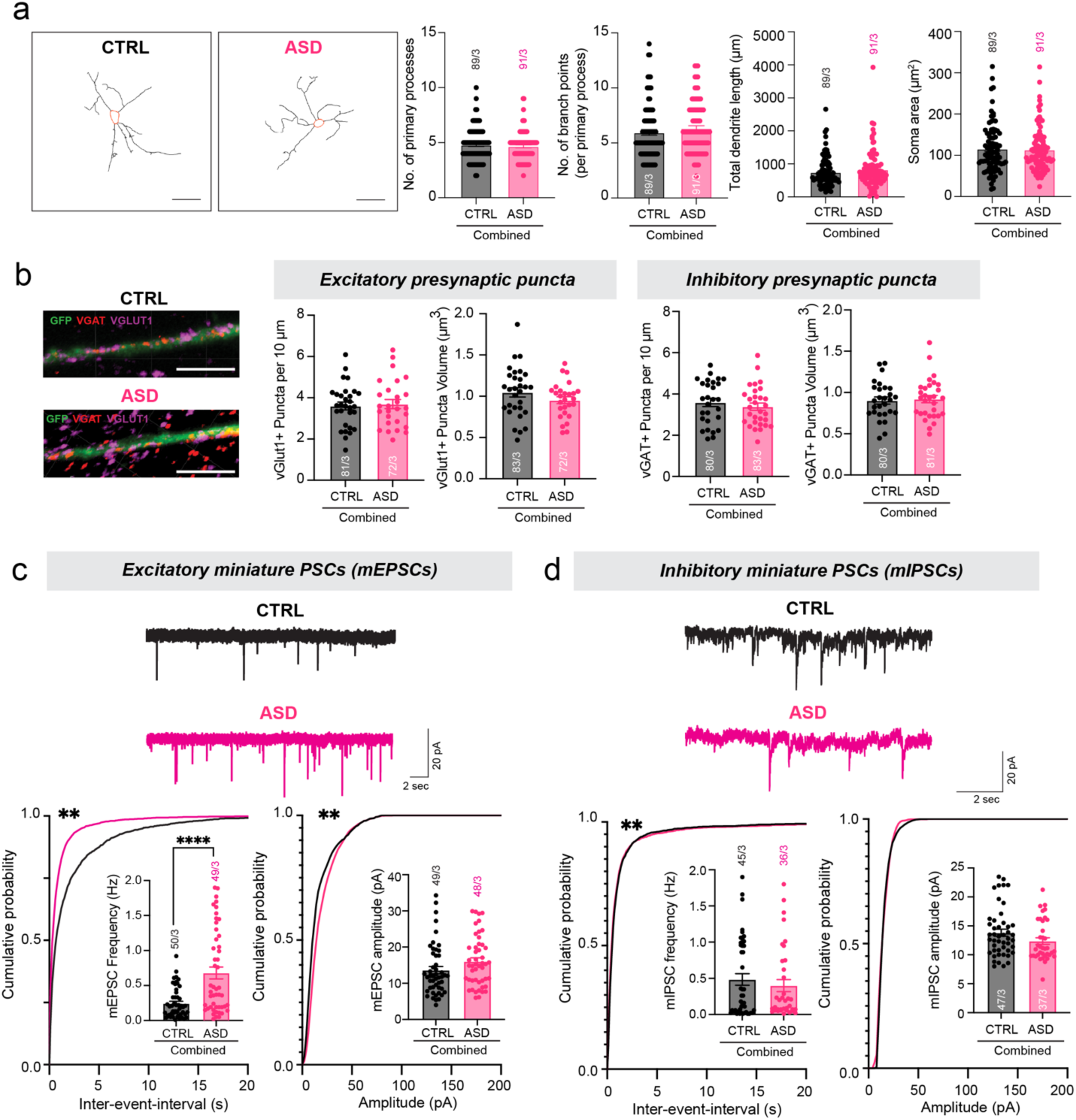
Increased spontaneous synaptic signaling in *NRXN1*^+/-^ ASD patient E-I iNs is mediated by higher frequency of excitatory synaptic vesicular release without changes in synapse formation and neuronal morphology. a) Left: representative binary mask of EGFP+ excitatory iNs in the E-I mixed cultures (scale bar, 40 µm); Right: quantification summary graphs of various parameters of neuronal morphology (number of primary processes, number of branch points per primary processes, total dendritic length, and soma size) in control vs. ASD iNs. **b)** Left: representative images of isolated EGFP+ excitatory neuronal dendritic segments stained for vGlut1 (excitatory synapse marker) and vGAT (inhibitory synapse marker) (scale bar, 5 µm). Note Ngn2-iNs in the E-I mixed cultures are specifically labeled with EGFP due to the presence of TetO-EGFP in these neurons (see Fig. 1); *Right panel*: Quantification summary graphs of the puncta density and volume in control vs. ASD iNs. **c)** Top: representative spontaneous miniature postsynaptic excitatory current (mEPSC) traces; Bottom: pooled summary graphs of *NRXN1*^+/-^ ASD patient and control recordings of mEPSC frequency (left) and amplitude (right). Recordings were performed in the presence of TTX (1 uM) to block all APs to capture single vesicular release miniature events and in PTX (50 uM) to inhibit GABAergic inhibitory postsynaptic currents. Quantifications are shown both as cumulative probability plots and as bar graphs (inserts). **d)** Top: representative spontaneous miniature postsynaptic inhibitory current (mIPSC) traces; Bottom: pooled summary graphs of *NRXN1*^+/-^ ASD patient and control recordings of mIPSC frequency (left) and amplitude (right). Recordings were performed in the presence of TTX (1 uM) to block all APs to capture single vesicular release miniature events and in CNQX (20 µM)/APV (50 µM) to inhibit AMPAR- and NMDAR-mediated glutamatergic postsynaptic currents. Quantifications are shown both as cumulative probability plots and as bar graphs (inserts). All summary data for morphology are means +/- SEM (numbers in bars or parentheses show number of cells/independent cultures analyzed). Cells were fixed for analysis on dpi 49. No significant differences were detected in any of the conditions by Student’s t-test. Nonsignificant comparisons are not indicated. All summary data for minis are means +/- SEM (numbers in bars or parentheses show number of cells/independent cultures analyzed). Cells were recorded at 42- 49 dpi for mEPSCs and at 49-56 dpi for mIPSCs. Statistical significance for average frequency and amplitude were determined with Student’s T-Test, whereas significance for cumulative probability distributions was determined by two-tailed Kolmogorov-Smirnov test. P-values: p**< 0.01, p****<0.0001.

### Increased probability of excitatory neurotransmitter release underlies fast spontaneous synaptic signaling observed in NRXN1+/- ASD patient-derived E-I iNs

Increased frequency of ‘spontaneous synaptic events can be contributed by either increased neurotransmitter release probability or increased synapse numbers presynaptically. Given that increased synapse density is ruled out, we tested miniature synaptic events (AP-independent spontaneous single vesicular fusion events) by recording pharmacologically isolated spontaneous miniature excitatory (mESPCs) and miniature inhibitory (mIPSCs) postsynaptic current amplitude and frequency. Changes in mini frequency correlate with either a change in neurotransmitter release probability or synapse number whereas mini amplitude changes correspond to either a change in quantal size or postsynaptic receptor composition^29,30,31^. We observed a significant increase in the frequency of mEPSCs without changes in the mEPSC amplitude in ASD patient iNs compared to controls (**Figure 4c**). Interestingly, ASD patient iNs did not exhibit differences in mIPSC frequency or amplitude (**Figure 4d**), suggesting that GABAergic transmission was intact. This effect was consistent even when ASD patient iNs were analyzed in individual pairs (**Figures S4, S5**) with some variability across pairs noted.

The specific change in excitatory mEPSC frequency without affecting GABAergic mIPSCs implicates that E-I balance could be shifted primarily through increased excitation via increased probability of glutamate release. To test this idea, we performed evoked synaptic release recordings by delivering two sequential stimuli with varying stimulation intervals (paired pulse stimulations). By calculating the ratio between the second and first evoked EPSC amplitudes, we compared the paired pulse ratios (PPR) between a specific pair of ASD patient and control iNs (pair 1; **Figure 5b**). ASD patient iNs displayed lower PPR compared to control iNs, especially at the 25 ms stimulus interval, indicating increased initial release probability. By averaging the first evoked EPSC amplitudes, we found a trending increase in excitatory synaptic strength in ASD patient iNs as well (**Figure 5a**, p=0.07). This specific synaptic phenotype is completely opposite of what we have previously reported for *NRXN1*+/- SCZ and cKO iNs^16^, which showed increased PPR and decreased initial neurotransmitter release probability. Altogether, our results from mini and evoked recordings indicate that *NRXN1*+/- ASD iNs exhibit enhanced excitatory synaptic signaling through enhanced presynaptic glutamate release, which may, in turn, disrupt E-I balance and synaptic homeostasis.

**Figure 5.**
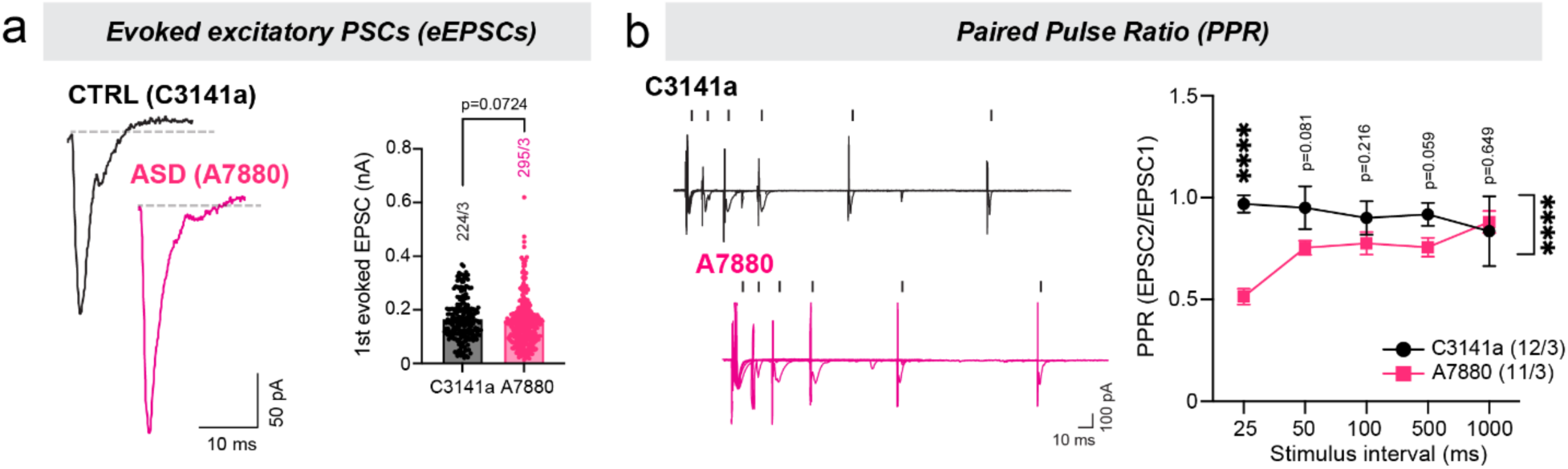
*NRXN1*^+/-^ ASD patient E-I iNs increase the initial probability of excitatory neurotransmitter release. **a)** Left: representative traces of evoked EPSCs; Right: summary graph of the EPSC amplitudes. **b)** Left: representative traces of eEPSCs recorded in response to two sequential action potentials elicited with 25- to 1000-ms intervals; Right: Summary plot of the PPR of EPSCs elicited with 25- to 1000-ms intervals showing decreased PPR primarily at the 25- ms interval. Summary data for 1^st^ EPSC amplitudes are means +/- SEM (numbers in bars or parentheses show number of 1^st^ evoked responses/2 batches analyzed). Cells were recorded at 49 dpi. Statistical significance for comparisons of average amplitudes across genotypes at each stimulus interval (for both 1^st^ evoked and PPR) was performed with Student’s T-Test and the genotype comparisons using two-way ANOVA with Tukey’s post hoc test. P-values are indicated at each stimulus interval and P-values: p****<0.0001.

### Functional connectivity analysis reveals irregular excitatory neuronal networks in NRXN1+/- ASD patient-derived E-iNs

Given that changes in the excitatory synaptic signaling were primarily eliciting the resulting synaptic phenotypes in *NRXN1*+/- ASD iNs, we switched to Ngn2-iN solo cultures for high-density microelectrode array (HD-MEA) recordings. In a previous study, we have shown that Ngn2-iNs display robust neuronal spiking and synchronized bursts, indicative of strongly connected neuronal networks^27,32^. Using the same experimental paradigm, we performed HD-MEA recordings from all three pairs of ASD patient and control iNs (E-iNs specifically). Recorded neuronal spikes were sensitive to synaptic blockers indicating that they are synaptically driven responses (**Figure S6**). To better correlate the data obtained from HD-MEAs with data from synaptic recordings (patch clamp), we subjected the neuronal spike data to a spike sorting algorithm (Kilosort)^33^ and further extracted functional connectivity maps using spike train cross- correlation analysis^34^. Using the Poisson distribution estimation of the peak in the cross-correlogram (CCG)^34^, functionally connected units were labeled as sender, receiver, or relay based on signal direction (**Figure 6a**). Signal strength between units or spike transmission probability between a sender-receiver pair was then calculated using the difference between the CCG peak and the baseline in the same bin, divided by the number of pre-synaptic spikes (**Figure 6b**). Using this analysis method, we found that ASD patient E-iNs showed a trend toward enhanced transmission probability (**Figures 6c**, **S7a**), as well as a statistically significant increase in the coefficient of variation (CV) of inter-spike-intervals (ISI) of the receivers (**Figures 6d**, **S7b**), indicating presence of more frequent irregular firing patterns. When examining the normalized distributions of ISIs, similar distributions were apparent between genotypes (**Figure S7c**), suggesting specific alterations in connectivity patterns of *NRXN1*+/- ASD patient E-iNs compared to controls.

**Figure 6.**
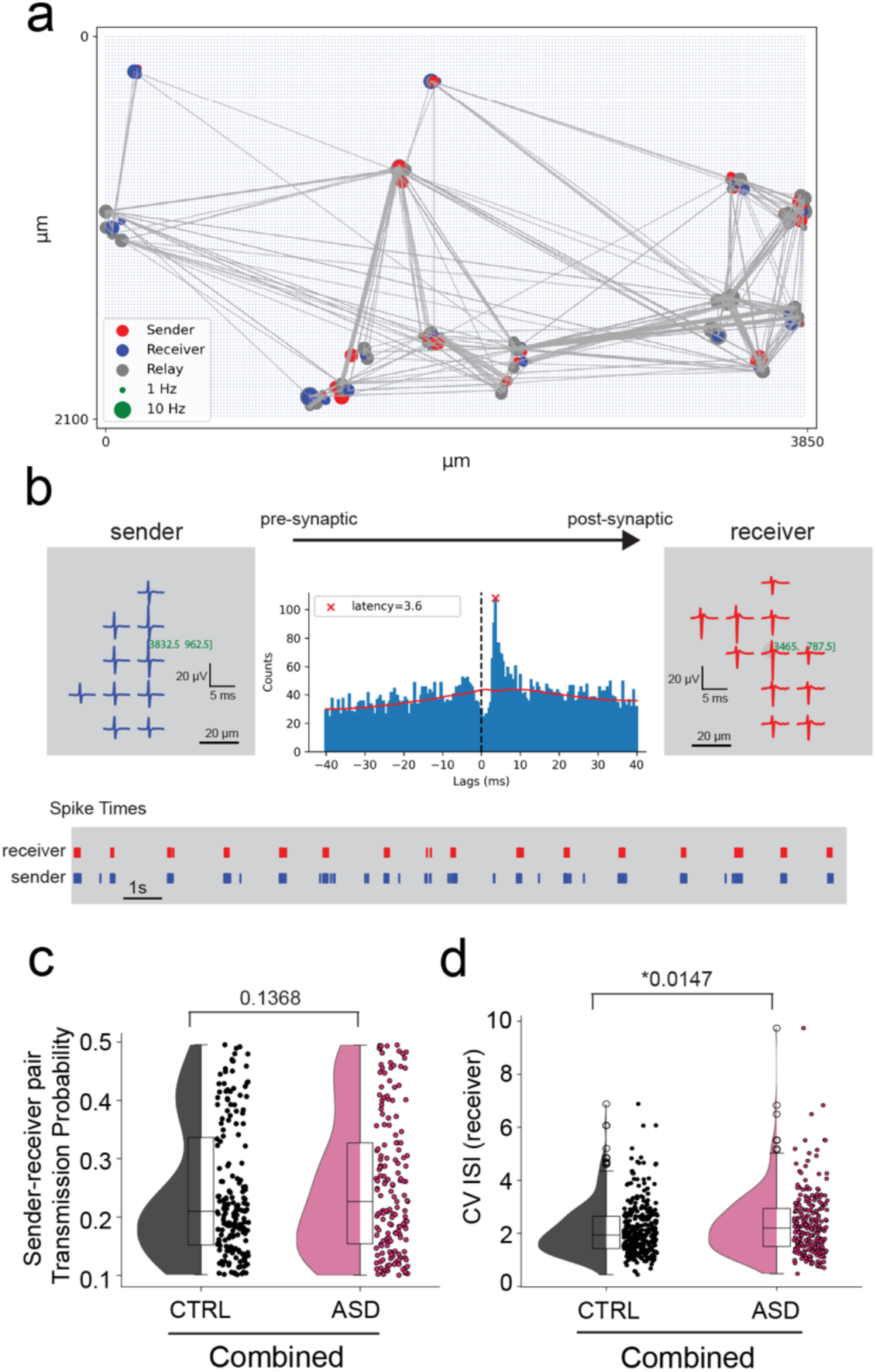
Functional connectivity analysis shows trending enhancement in connectivity probability and significant irregular firing patterns in excitatory neuronal networks from NRXN1+/- ASD patient iPSCs. HD-MEA recordings obtained from 28 dpi Ngn2 solo cultures from 3 *NRXN1^+/-^* ASD vs. 3 control iPSC lines (n>6 independent cultures per line). **a)** Units displaying functional connectivity across distances on the HD-MEA. Units are spike sorted, and functional connectivity is determined using spike train cross-correlation. A unit that only sends action potentials is labeled a sender. A unit that only receives action potential from other units is labeled a receiver. A unit that both sends and receives action potential is labeled a relay. Different colors are used on the functional connectivity map to distinguish these roles (sender-red, receiver-blue, and relay-gray). Lines are drawn between functional unit pairs on the electrode map, with the line width corresponding to the spike transmission probability. **b)** Example of a pre-synaptic (sender) and post-synaptic (receiver) connection detected between two units with distinct footprints and firing patterns. The cross-correlogram indicates a 3.6 ms signal latency from receiver to sender (28 dpi Ngn2-iN A4927). **c)** Pre- and post-synaptic spike transmission probability in *Ngn2* cultures, with pooled analysis of the three control and three ASD genotypes (combined). Box plots showing median values of transmission probability with upper quartile (75%) and lower quartile (25%). Increased transmission probability indicates increased likelihood that sender and receiver are functionally connected. **d)** Coefficient of variation (CV) of interspike intervals (ISI) for receivers in *Ngn2* cultures, with pooled analysis of the three control and three ASD genotypes (combined). Box plots showing median values of CV ISI with upper quartile (75%) and lower quartile (25%). Increased CV ISI indicates more irregular firing patterns.

### Decreased excitatory and inhibitory synaptic transmission in NRXN1+/- SCZ patient and cKO E-I iNs

So far, findings in the mixed E-I iNs from *NRXN1*+/- ASD patient iPSCs all point towards hyperconnectivity, a phenotype which we have yet to observe in any previously studied *NRXN1* deletion iNs. Therefore, we differentiated E-I iNs from two independent *NRXN1*+/- SCZ patient iPSCs with varying deletion sizes (N3320a and N1884a^16^) and performed mEPSC and mIPSC analysis. As expected, *NRXN1*+/- SCZ iNs showed decreased synaptic transmission overall, resulting in decreased mEPSC frequency and amplitude as well as decreased mIPSC frequency for the N3320a line (**Figure 7**). An independent experimental replication was performed using a second SCZ patient line N1884a, which also displayed decreased mEPSC and mIPSC frequencies specifically (**Figure S8**). Notably, the N1884a patient line carries a similar deletion size and breakpoint as two of the ASD patients whereas N3320a patient carries a larger deletion in the *NRXN1* locus (**Figure 1b**). Finally, to confirm the decreased synaptic transmission phenotype observed in *NRXN1*+/- SCZ iNs, we differentiated E-I iNs from the *NRXN1* heterozygous cKO line (exon 19 deletion)^17^ and performed mEPSC and mIPSCs recordings. *NRXN1* cKO iNs also displayed decreased mEPSC and mIPSC frequencies like SCZ patient iNs (**Figure S9**), further validating dampened synaptic signaling in cKO.

**Figure 7.**
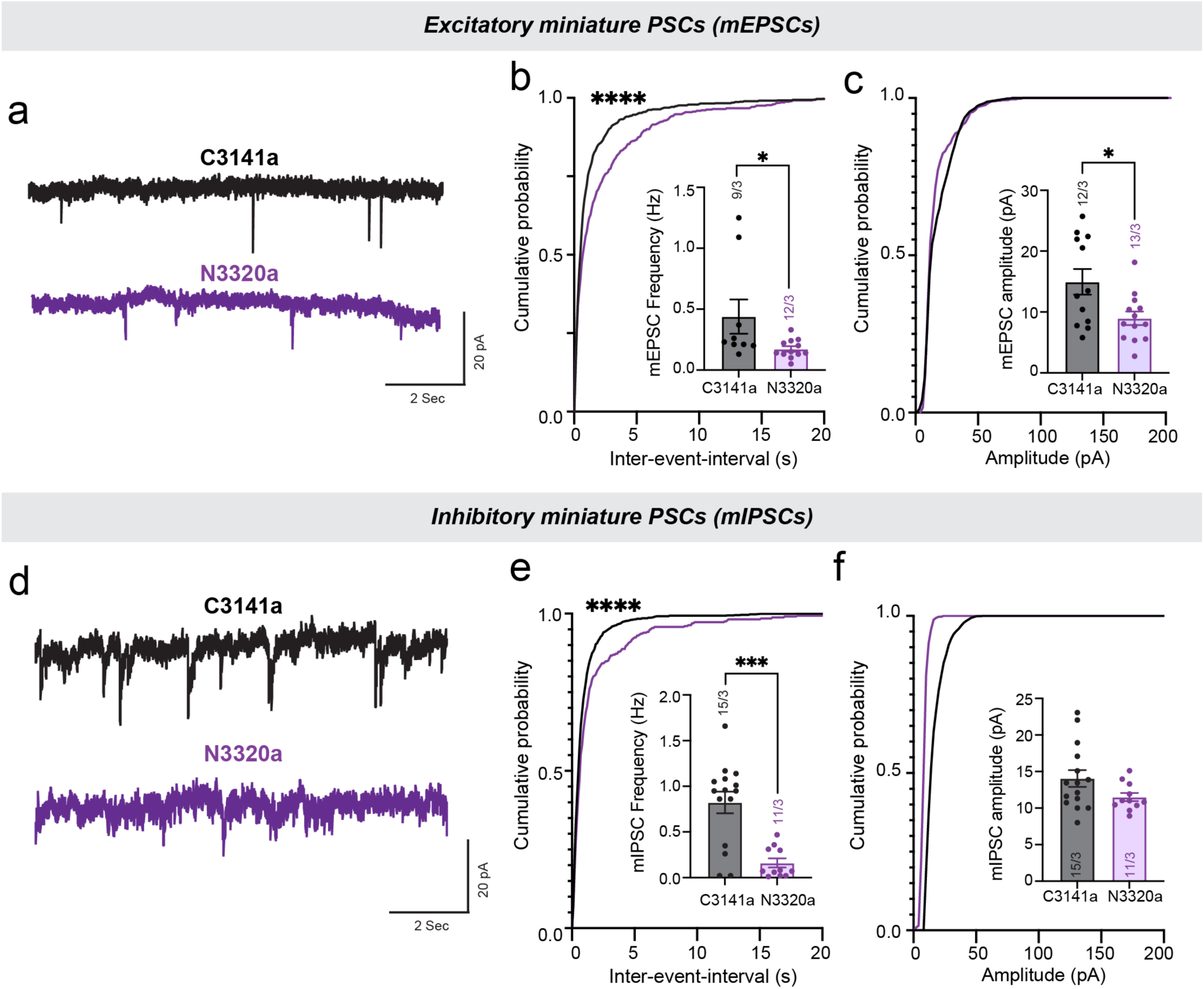
*NRXN1^+/-^*SCZ patient-derived E-I iNs show decreased excitatory and inhibitory synaptic transmission, opposite of ASD phenotypes. Same E-I iN culturing scheme and patch recording of EGFP-labeled excitatory iNs were applied to C3141a and N3320a as described in Figure 1. Recordings were performed in the presence of TTX (1 µM) to block all APs to capture single vesicular release miniature events and in 1) PTX (50 µM) to inhibit all GABAergic inhibitory postsynaptic currents for mEPSCs; and 2) CNQX (20 µM)/APV (50 µM) to inhibit AMPAR- and NMDAR-mediated glutamatergic postsynaptic currents for mIPSCs. Quantifications are shown both as cumulative probability plots and as bar graphs (inserts). **a)** Representative mEPSC traces and quantifications of mEPSC frequency **(b)** and amplitude **(c). d)** Representative mIPSC traces and quantifications of mIPSC frequency **(e)** and amplitude **(f).** All summary data are means +/- SEM (numbers in bars or parentheses show number of cells/independent cultures analyzed). Cells were recorded at 42-49 dpi for mEPSCs and 49-56 dpi for mIPSCs. Statistical significance for average frequency and amplitude were determined with Student’s T-Test, whereas significance for cumulative probability distributions was determined by two-tailed Kolmogorov-Smirnov test. P-values: *p< 0.05, ***p<0.001, ****p<0.0001.

### Differential effects on homeostatic synaptic upscaling in NRXN1^+/-^ ASD vs. SCZ patient-derived E-I iNs

Homeostatic synaptic plasticity is a key homeostatic mechanism which ensures balanced and stable activity levels in response to changes in neuronal activity in the developing cortical networks^31,35,36^. A specific type of homeostatic synaptic plasticity called synaptic upscaling is a rebound phenomenon occurring in response to chronic neuronal silencing and has been observed mouse models both *in vitro* and *in vivo*^31,35–37^. Interestingly, synaptic scaling has been shown to be disrupted in ASD-associated single gene disruptions in mouse models of *Fmr1* and *Shank3* KO^37,38^. We reasoned that, since *NRXN1*+/- ASD iNs show altered balance in E-I synaptic signaling, *NRXN1*+/- ASD iNs may exhibit disrupted synaptic homeostasis. Application of TTX in control E-I iN cultures for 48 hrs triggered expected synaptic upscaling as evidenced by increased sPSC amplitudes, an indication of increased synaptic strength (**Figure 8** – see C8905 and C3141a). In contrast, *NRXN1*+/- ASD patient iNs could not trigger synaptic upscaling like control counterparts and unexpectedly rescued its enhanced sPSC amplitude response (**Figures 8a,b**).

**Figure 8.**
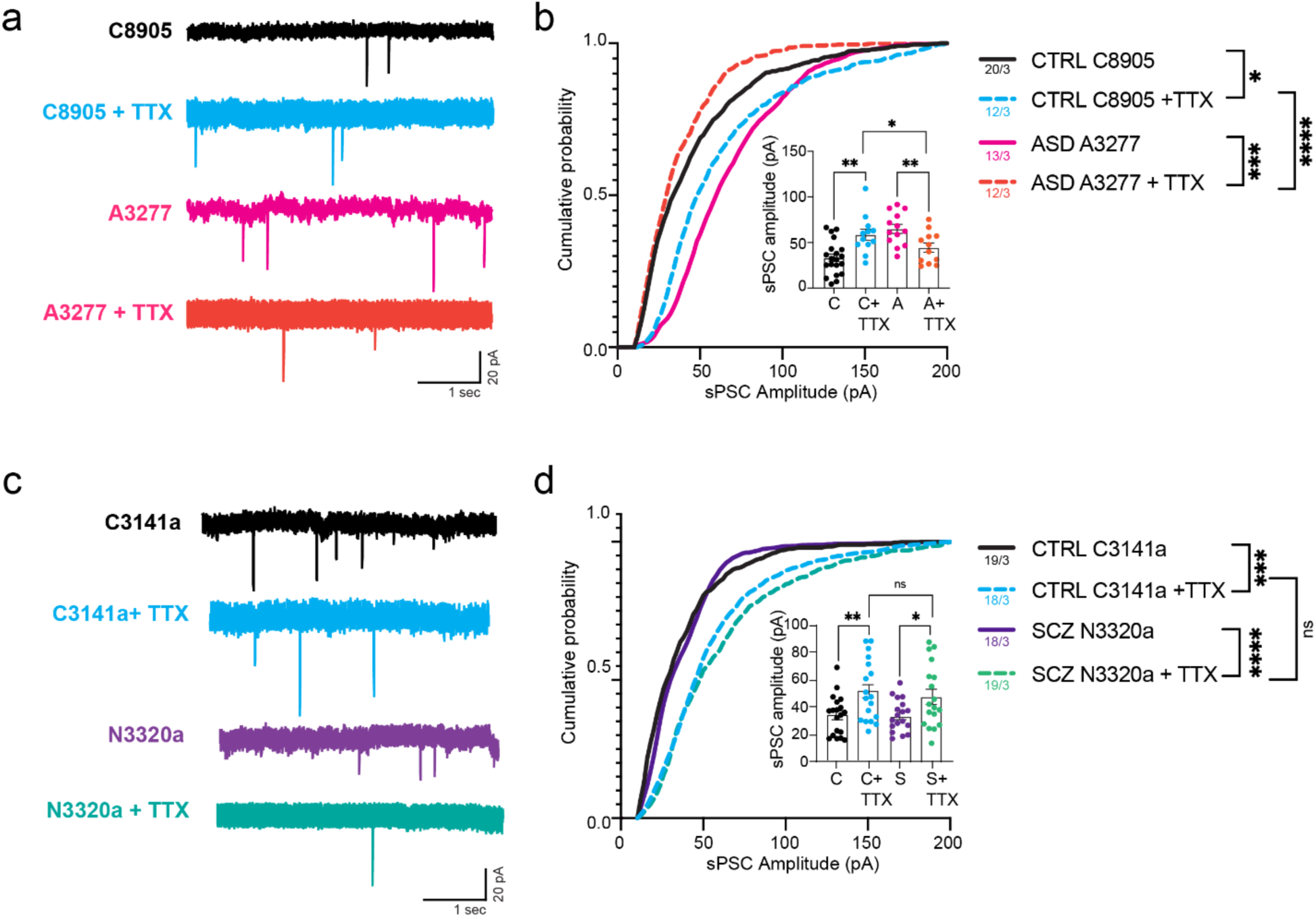
Differential effects on homeostatic synaptic upscaling in *NRXN1^+/-^* ASD vs. SCZ patient-derived E-I iNs suggest disorder-specific synaptic plasticity deficits. TTX-mediated homeostatic synaptic upscaling is impaired in *NRXN1^+/-^* ASD E-I iNs but intact in *NRXN1^+/-^*SCZ E-I iNs. Representative traces of sPSCs recorded from E-I iN control vs. *NRXN1^+/-^* ASD (pair 2) **(a)** as well as E-I iN control vs. *NRXN1^+/-^* SCZ (N3320a) **(c)** with or without chronic neuronal silencing. E-I iNs were treated for 48 hrs with TTX (1 µM), followed by a 30-min washout before recording. **b, d)** Analysis of sPSC amplitudes of control, ASD and SCZ lines with or without TTX treatments. Quantifications are shown both as cumulative probability plots and as bar graphs (inserts). All summary data are means +/- SEM (numbers in bars or parentheses show number of cells/independent cultures analyzed). Cells were recorded a 49-56 dpi for sPSCs. Statistical significance for average amplitude were determined with Student’s T-Test within each genotype conditions (C vs. C+TTX or A or S vs. A+TTX or S+TTX), whereas significance for cumulative probability distributions was determined by two-tailed Kolmogorov-Smirnov test. P-values: *p< 0.05, p**< 0.01, ***p<0.001, ****p<0.0005.

Intriguingly, synaptic upscaling was intact when the same experimental paradigm was tested in *NRXN1*+/- SCZ patient iNs (**Figures 8c,d**), suggesting that homeostatic synaptic plasticity is uniquely impacted in the ASD patient background.

## Discussion

In this study, we performed comprehensive functional analyses on a new cohort of *NRXN1*+/- ASD iPSCs by differentiating patient iPSCs into E-I iNs, a mixed co-culture scheme that allows functional dissection of E-I signaling, network balance, and synaptic homeostasis. These patient iPSC-iNs were paired randomly with well characterized healthy control iPSC-iNs^16^ for synchronized culturing and downstream assays (**Figure 1**). Notably, we found that *NRXN1*+/- ASD E-I iNs display enhanced synaptic connectivity, as demonstrated by more frequent fast spontaneous synaptic events (IEI analysis of sPSCs; **Figure 3**), an increased frequency of excitatory miniature synaptic release (mEPSCs; **Figure 4**), enhanced excitatory synaptic strength in excitatory synapses (eEPSCs; **Figure 5a**) accompanied by a higher release probability (decreased PPR; **Figure 5b**). These parameters of basal synaptic transmission are altered in the absence of changes in the intrinsic membrane properties, AP properties, and neuronal excitability (**Figure 2**). In addition, overall dendritic morphology and synapse numbers were unchanged between *NRXN1*+/- ASD patient iNs and controls (**Figure 4 a,b**). Such results are contrary to what was previously reported in the *NRXN1*+/- SCZ patient and *NRXN1* cKO excitatory iNs^16,17^. To further confirm opposing synaptic phenotypes in a mixed E-I iN culture setting, we performed electrophysiological recordings in two independent *NRXN1*+/- SCZ patient lines and a *NRXN1* cKO line (exon 19 heterozygous deletion) (**Figures 7**, **S8, S9**). Both *NRXN1*+/- SCZ patient iNs and *NRXN1* cKO iNs showed an overall decrease in excitatory and inhibitory synaptic transmission, further validating weakened synaptic transmission and decreased neurotransmitter probabilities seen in these models^16,17^.

Increased excitatory synaptic transmission seen in *NRXN1*+/- ASD patient iNs translated to irregular neuronal firing patterns in the network (**Figure 6**) and impaired homeostatic synaptic plasticity (**Figure 8**). Intriguingly, synaptic scaling after chronic activity silencing with TTX treatment was not affected in *NRXN1*+/- SCZ patient iNs (**Figure 8**). It may be possible that such differences in synaptic homeostasis could be explained by their overall basal synaptic transmission states. In ASD patient iNs, excitatory synaptic drive could be already at its maximum, and therefore, homeostatic synaptic upscaling cannot be further induced (occlusion). In SCZ patient iNs, synaptic upscaling is normal. However, cortical organoids derived from these patient lines could not potentiate their synchronized network responses in a chemical LTP paradigm, mimicking another form of synaptic plasticity^18^. Therefore, it is tempting to postulate that, although synaptic upscaling is intact in *NRXN1*+/- SCZ patient iNs, synaptic downscaling (Bicuculine induced) could be altered due to their dampened basal synaptic states. Nonetheless, the distinct ways in which two different disorder patient lines respond to synaptic plasticity paradigms are interesting and warrants further future studies dissecting which exact molecular and cellular pathways are differentially impacted.

Interestingly, an ASD-associated mutation in the *NLGN3* gene, a postsynaptic ligand of NRXN1, induces hyperconnectivity in a similar mixed E-I iN culture model^26^. Notably, the aberrant synaptic deficits observed in *NRXN1*+/- ASD patient iNs closely resemble those caused by the *NLGN3* R451C gain-of-function mutation linked to ASD^26^. Functionally, *NLGN3* R451C iNs exhibit increased excitatory synaptic strength without altering inhibitory synaptic signaling^26^, a pattern also seen in *NRXN1*+/- ASD patient iNs. These findings suggest that both a single ASD-associated variant and the broader ASD patient genetic background converge on the same excitatory synaptic augmentation, further reinforcing the E-I imbalance hypothesis in ASD^39^.

There are several implications for the present study. First, based on our work here as well as prior reports^16,17^, *NRXN1*+/- deletions in the patient populations will not consistently lead to the exact same synaptic phenotypes but rather will depend on their disorder background. We showed that, while *NRXN1*+/- ASD iNs produce enhanced excitatory synaptic signaling leading to overall hyperconnectivity, *NRXN1*+/- SCZ patient iNs exhibit weakened excitatory/inhibitory signaling leading to hypoconnectivity. This argument is supported by the fact that iNs from SCZ patient (N1884a) and ASD patients (A3277, A7880) who carry similar genomic deletion location and size produce opposite phenotypes. Most likely, the polygenic burden specific to the disorder together with *NRXN1*+/- deletions will dictate the final cellular outcomes. Our findings suggest a different interpretation from a recent study by Fernando et al.^40^, wherein opposing functional phenotypes are reported in *NRXN1*+/- deletion patient iPSCs with 5’ and 3’ deletion locations. Their reported 5’ deletion carrier with bipolar disorder diagnosis (BP) present comparable copy number deletion size and location to our SCZ and ASD patient lines (N1884, A3277, A7880; **Figure 1b**). Yet, the BP patient iNs exhibit weakened excitatory synaptic signaling, mimicking only the SCZ patient cases but not ASD patient cases in our analyses. Interestingly, iNs from their 3’ deletion carrier with SCZ diagnosis show enhanced excitatory synaptic signaling, mirroring that of our ASD patient cases. In the future, it will be informative to examine and compare full length transcript isoforms generated from the *NRXN1* locus in all of these patient lines to further test the functional contribution of mutant isoforms in neurophysiology.

Second, the differential homeostatic plasticity phenotypes observed in SCZ vs. ASD patient iNs suggest that each disorder may exhibit unique compensatory adaptations in network properties in response to altered neuronal activity. We speculate that such adaptive mechanisms may underlie differential onset of symptoms and clinical manifestations. For example, SCZ is a later onset disorder, peaking at late adolescence, whereas ASD is an early onset, appearing typically within first two years of birth. In SCZ, hypofunctioning of glutamatergic synapses is a well-accepted hypothesis^41^ and is in line with synaptic pruning hypothesis where over-pruning of weakened synaptic connections is thought to be one of the key pathological mechanisms^42^. In ASD, hyperconnectivity and hyperexcitability seem to be the predominant neurophysiological mechanisms in humans and *in vitro* stem cell-based preclinical models^43–47^. Though it is extremely difficult to separate the initial causal physiological mechanism from homeostatic compensation, testing how patient derived neurons exhibit plasticity in various perturbations and activity manipulations will inform network- and synapse-level signaling dynamics that may be differentially altered in one disorder over the other.

Although this work provides valuable insights into *NRXN1* variant mediated pathogenesis in two different disorder contexts, certain limitations exist. First, because iNs bypassed developmental stages, the early neurodevelopmental influences impacting initial synapse formation are not accounted for. As ASD especially exhibit an early neurodevelopmental component^48^, in the future, it will be important to utilize complex 3D neural organoid models to examine how early neurodevelopmental processes impact circuit development *in utero* and how the diseased circuits become modified and remodeled post-birth. Second, because there is no perfect ‘biological control’ that can be paired with the patient iPSC lines bearing large deletions in the *NRXN1* locus, our study utilized sex- and passage-matched healthy control iPSCs. Due to this limitation, we chose to report trends and patterns that emerged from all three ASD patient iPSCs compared to all three control iPSCs. At the individual pair level, there certainly is a degree of variation across lines (see individual pair analyses in **Figures S1,S4,S6,S7**). This level of variability is expected given that each individual iPSC line is unique and have variable genetic backgrounds. Third, the rare nature of *NRXN1* deletions makes it difficult to obtain and screen through a large cohort of iPSC lines from patients. Eventually, the scope of the cell lines needs to be further expanded in the future to capture the genetic and biological complexities and heterogeneity associated with *NRXN1* deletions. Here, all iPSC lines used are from Caucasian males and since sex and ancestry may influence the penetrance and clinical expressivity of *NRXN1* deletions^9,49^, expanding the cell line collection to include females and other ancestry cohorts will be important. Finally, inclusion of neurotypical individuals who are unaffected phenotypically by *NRXN1* deletions will reveal molecular markers of neuroprotective mechanisms and/or resilience.

## Methods

### Human iPSC generation

Human iPSC cell lines (SAMPLED ID’s SSC07880, SSC04927, SV0003277) were established by the Simons Foundation Autism Research Initiative (SFARI). These lines each come from a male patient diagnosed with an autism spectrum disorder and carry the following genomic changes at 2p16.3 - SSC07880 (SFARI ID 13580-p1) harbors a 64 kb deletion at Chr2: 51,020,156- 51,084,394; SSC04927 (SFARI ID 12501-p1) harbors an A-to-T substitution at Chr2: 50,497,467, a frame shift point mutation resulting in an early termination (Tyr955Ter); SV0003277 (SFARI ID 17980-X2) harbors an 88 kb deletion at Chr2: 51,018,339-51,106,818. All genomic coordinates provided utilize the hg38 assembly. Our line ID uses the last four digits of each line, i.e. A7880 = SSC07880, A4927 = SSC04927, A3277 = SV0003277. All human iPSC lines have been tested normal karyotype and negative for mycoplasma.

Two individuals (A7880, A4927) were part of the Simons Simplex Collection (SSC) and one individual (A3277) came from Simons Searchlight cohort. We are grateful to all of the families at the participating Simons Searchlight sites and Simplex Collection (SSC) sites, as well as the principal investigators involved in the collections. Approved researchers can obtain the biospecimens used in this study by applying at https://base.sfari.org.

### Stem Cell Culture

Human iPSCs/ESCs were cultured and maintained as feeder-free in MTeSR Plus medium (Stem Cell Technologies) on Matrigel (Corning)-coated plates in 37°C, 5% CO_2_. In brief, human iPSCs were maintained with daily media changes, and human ESCs were maintained with media changes every other day. Cells were split upon 70% confluency with ReLeSR (Stem Cell Technologies) and passaged in the presence of Y-27632 (10 µM, Axon Medchem).

### Lentivirus Generation

Lentiviral constructs containing Tet-On versions of *Ngn2, Ascl1, Dlx2,* and *EGFP*, as well as *rtTA, FLP, CRE, and TdTomato* were generated in HEK293T cells (ATCC) as previously described^17,23,25^. In brief, HEK cells were grown to 80-90% confluency in T75 flasks in DMEM containing 10% FBS (Sigma) and 1% Penn-Strep. Upon confluency, a fresh media change with exactly 12 mL was performed before calcium phosphate transfection of the cells with 12 µg of the appropriate inducible gene-containing plasmid as well as the following viral helper plasmids: 8.1 µg pMDLg/pRRE, 3.9 µg of pRSV-rev, and 6 µg of VSVG. DNA was mixed in 900 uL of ultrapure water containing 115 µL of 2.5M CaCl_2_. This was added dropwise while gently vortexing to 900 µL 2x HBS (280 mM NaCl, 1.5 mM Na_2_HPO_4_, 50 mM HEPES adjusted to pH 7.1). The transfection mixture was incubated at room temperature for 20 minutes before being added dropwise to the cells. Cells were then incubated for 5 hours at 37°C and 5% CO_2_, then media was changed. After 48 hours, the media was collected, pelleted at 300xg for 5 minutes, and the supernatant was collected and filtered through a 0.45 µM cellulose acetate filter into an ultracentrifuge tube. This was pelleted at 61,500 xg for 2 hours at 4°C. Supernatant was completely removed, and the pellet was resuspended in 100 µL MEM and aliquoted for use.

### Generation of excitatory and inhibitory human neurons

NGN2 and A/D cells were differentiated as previously described^23,50,25^. In brief, stem cells were lifted with Accutase (Innovative Cell Technologies) and counted with a Countess 3 automated cell counter with trypan blue. For 1 million cells, 10 µL of *Ngn2,* rtTA, and TetO-EGFP lentiviruses were added to 4 mL mTeSR plus with Y-27632 along with the cells into a 15 mL conical tube and incubated for 5 minutes at room temperature. The volume was then adjusted to 12 mL with media and plated on Matrigel-coated plates. The same process was used for A/D cells, except for 1 million cells, 20 µL rtTA, *ASCL1,* and DLX2 Lentiviruses were used and for indicated experiments, 10 µL of TetO-TdTomato were also added. For *NRXN1* het cKO line, 10 µL of either FLP or CRE lentivirus was also added to the infection mixture with TdTomato without TetO-EGFP. The cells were allowed to expand for 1-2 days in mTeSR plus media before being induced in *induction media* (DMEM/F12 (ThermoFisher) with 1X Non-Essential Amino Acids (Gibco), 4 µg/mL Doxycycline (Clonetech), 1X N2 supplement (ThermoFisher), 10 ng/mL Brain-Derived Neurotrophic Factor (BDNF; Peprotech), 200 ng/mL mouse laminin (Life Technologies), and 10 ng/mL Neurotrophin-3 (NT3; Peprotech)). This is considered day 0 post induction (dpi). 1-4 dpi, media was replaced with *selection media* (*induction media* with the appropriate selective antibiotics added; 10 µg/mL puromycin (Invivogen) and 300 µg/mL hygromycin-B (Thermo Fisher)). At 5 dpi, cells were lifted with Accutase (Innovative Cell Technologies) and a cell lifter as necessary, pelleted at 300 xg for 5 minutes at room temperature, and resuspended in Bambanker freezing media (Wako Chemical) and frozen in vials at 2-4 million cells per vial.

### Mouse Glia culture

Newborn pups (p0-p2) were harvested and sacrificed by decapitation as previously described^27^ and in accordance with University of Massachusetts Amherst IACUC protocol #5953. Briefly, pup heads were placed in a dish containing 1X Hank’s Balanced Salt Solution (HBSS, ThermoFisher) and the brain was removed. The meninges were carefully removed, along with the olfactory bulbs and cerebellum, leaving only the outer cortex and hippocampal formation. Tissue was dissociated with a papain mixture (6.4 units/mL papain (Worthington Biochemical) and 500 µM EDTA in 1X HBSS) at 37°C for 20 minutes with occasional agitation. After incubation, clumps were decellularized with a 1 mL pipettor, washed with HEK media, and plated on T75 dishes. Media was changed the following day, and cells were split twice to increase yield and ensure no neuronal carryover. Glia were cryopreserved at 2 million cells per vial in freezing media (DMEM with 20% FBS and 10 % DMSO).

### Neuronal culture

Induced neurons were thawed and plated in *plating media* (Neurobasal (Gibco) supplemented with 1X Glutamax (Fisher), 1X B27 with vitamin A (Gibco), 4 µg/mL doxycycline, 10 ng/mL BDNF, 10 ng/mL NT3, 10 ng/mL Glial Cell Derived Neurotrophic Factor (GDNF; Fisher), and 200 ng/mL mouse laminin). Cells were counted with trypan blue, and for all experiments except for HD-MEA, 150,000 glia, 120,000 Ngn2, and 30,000 A/D cells were plated on each Matrigel-coated coverslip. For HD-MEA and evoked recordings, this same ratio was kept, but all numbers were doubled (for high-density). Pure excitatory cultures on mouse glia were also plated for some experiments, with 150,000 glia and 150,000 Ngn2 being plated per coverslip. Cells were maintained with ½ media changes every other day (*plating media* containing 2 µM Cytosine β-D-Arabinofuranoside Hydrochloride (AraC; Sigma)) until 14 dpi, at which point ½ media changes twice a week began with *maturation media* (MEM with 5% FBS (Hyclone), 1X B27 supplement, 2.5% v/v 20% glucose solution, 0.25% v/v 8% NaHCO3, 0.1 mg/mL transferrin (Gemini Bio), 500 µM L-glutamine (Gibco), and 2 µM AraC) until experiments.

### High Density Multi-Electrode Array Recordings

MaxOne chips (Maxwell Biosystems) were prepared and plated as previously described^27,32^. Briefly, chips were treated overnight with 1% Tergazyme (Sigma), then washed three times in DI water. Chips were sterilized by soaking in 70% ethanol for 30 minutes the brought into the biosafety cabinet and dried. 50 µL of 0.07% PEI (Sigma) was added directly to the electrode array and incubated at 37°C and 5% CO_2_ for one hour. PEI was then removed and replaced with 50 µL Matrigel and incubated at 37°C and 5% CO_2_ for another hour, after which point the Matrigel was removed and cells were plated as described above in 50 µL of *plating media* and incubated overnight at 37°C and 5% CO_2_ in a bioassay dish with a well of sterile DI water to prevent evaporation. The next day, 500 µL of *plating media* was added to the chip, and maintenance of cultures continued as described in the neuronal culture section. 4x sparse activity scans and 5 minute network scans were performed. For the network scan, 20 neuronal units with the 600 electrodes exhibiting the highest amplitudes were selected for recording.

### Spike sorting and auto-curation

HD-MEA recordings are spike-sorted into single unit activities using Kilosort2^33^ and auto-curated by quality metrics^51^. Recordings are processed using a cloud-based neuro-data pipeline^51^ for consistency and reproducibility. Before sorting, recordings are bandpass filtered with a frequency range of 300-6000 Hz. Auto-curation parameters are set to 5 RMS for signal-to-noise (SNR) ratio, 20% for interspike-interval (ISI) violation (Hill method^52^), and 0.1 Hz for firing rate. Units with a lower SNR, higher ISI violations, or lower firing rates are removed. The remaining units are saved with their spike trains, footprints, and locations on the HD-MEA.

Due to the HD-MEA’s high electrode density and small pitch, a neuron’s signal can be captured simultaneously by multiple electrodes. Footprints are defined as spikes recorded on a maximum of 12 neighbor channels, with an average spike waveform computed for each channel. Locations include the (x, y) coordinate for the best channel (defined as the channel that recorded the unit’s largest amplitude) and the coordinates of neighbor channels on the footprint. For each unit, the average spike waveform is calculated using up to 500 raw spikes with a waveform length of 2.5 ms (50 data points at a 20 kHz sampling rate).

### Functional connectivity analysis

Functional connectivity analysis is performed by spike train cross-correlation and Poisson distribution estimation of the peak in the cross-correlogram (CCG)^34^. Spike trains are converted into a sparse matrix using a bin width of 0.4 ms. CCGs are computed by cross-correlating each unit’s sparse spike train with all others over a time window of -40 ms to 40 ms. The CCG baseline is calculated using a 60% hollow Gaussian kernel with a sigma of 10. The coefficient for Poisson distribution estimation is derived from the CCG peak and its baseline. Units are considered functionally connected when the CCG peak fell between 1–5 ms and the coefficient was below 10^−5^. Spike transmission probability from a pre-synaptic neuron (sender) to a post-synaptic neuron is calculated using the difference between the CCG peak and the baseline in the same bin, divided by the number of pre-synaptic spikes.

### Whole-cell patch clamp electrophysiology

Cells were cultured as described above to 42 dpi, and through 56 dpi recordings were performed using Pclamp 10 software with an Axon Digitizer 1550b and a Multiclamp 700b amplifier (Molecular Devices). The digitizer was set to 10 KHz, while a 2 KHz low-pass filter was applied with the amplifier. For sPSCs, spontaneous action potential, voltage step, current step, and evoked recordings, cells were submerged in a bath solution (130 mM NaCl, 5 mM KCl, 2 mM CaCl, 1 mM MgCl_2_, 10 mM HEPES, and 10 mM glucose at pH 7.4 and 290 mOsm, osmolarity titrated with sucrose) and patched onto with glass pipettes pulled to have 5-9 MΩ resistances filled with potassium gluconate internal solution (126 mM K-Gluconate, 4 mM KCl, 10 mM HEPES, 4 mM ATP-Mg, 0.3 mM GTP-Na_2_, and 10 mM phosphocreatine at pH 7.2 and 290 mOsm). For miniature postsynaptic recordings the bath solution was supplemented with 1 µM TTX and the appropriate neurotoxins to isolate excitatory or inhibitory currents (50 µM picrotoxin for mEPSCs or 20 µM CNQX and 50 µM APV for mIPSCs) and the internal solution was cesium chloride-based (40 mM CsCl, 90 mM K-Gluconate, 1.8 mM NaCl, 1.7 mM MgCl2, 3.5 mM KCl, 0.05 mM EGTA, 10 mM HEPES, 2 mM ATP-Mg, 0.4 mM GTP-Na_2_, 10 mM phosphocreatine at pH 7.2 (with CsOH) and 290 mOsm). For evoked recordings, PTX (50 µM) was added to the bath to silence inhibitory signaling, and the internal solution was supplemented with QX-314 (5 mM, Tocris). Recordings were analyzed using Clampfit 11 software with in-house made templates for sPSCs, mEPSCs, and mIPSCs derived from each recording type. In voltage clamp protocols (sPSCs, minis, V-Step), cells were held at -65 mV throughout recording. For sodium and potassium current recordings (V-step), voltage was incrementally increased from -100 mV to +80 mV in 10 mV increments. For I-clamp recording (I-step), current was added until the cell hovered at around -63-65 mV before initiating recording, wherein current was applied to step the cell from -50 mV to +130 mV in 10 mV increments. For spontaneous action potential recordings, no holding was used. For evoked recordings, cells were primed with a −75 mV pattern before holding at −70 mV for the duration of the 20-second epoch (seven sweeps per recording). Pulse widths for paired-pulse stimulation were 25 ms, 50 ms, 100 ms, 500 ms, and 1000 ms, and all cells were stimulated with 1.9 mA of current directly over a group of at least 3-4 cells approximately 100 µm from the target cell.

### Electrophysiology analysis

The experimenter was blind to the genotype or cell line identity during recording and analysis, and were unblinded only after analysis was complete. Data organization and simple calculations such as the absolute amplitude (peak amplitude – baseline amplitude), Frequency (number of events / time of analysis window), and inter-event-interval (event start time – previous event start time) and binning for cumulative probability were performed here, then any values more than 2 standard deviations from the mean were excluded as outliers. The data were then exported to Prism 10.2 for further statistical analysis and graphing. For all metrics except for PPR and cumulative probability, a two-tailed Student’s T-Test was performed, and significance is represented by corresponding asterisks: *p<0.05, **p<0.01, ***p<0.001, ****p<0.0001; nonsignificant comparisons were not generally included. All P values can be found in the attached statistics worksheet.

For sPSCs: events were identified based on a template derived from these mixed cultures with clearly defined events. Events originating from bursts were manually excluded, and amplitudes below -200 pA were removed to ensure no bursts were included. Events above -10 pA were also excluded to ensure proper filtration of baseline noise.

For mini PSC recordings: a template specific for mEPSCs and mIPSCs were derived using the best recordings with clear signals for each event type. No bursts were observed. Events were excluded if they fell below -1 pA for mEPSCs and -8 pA for mIPSCs to minimize noise conflation.

For spontaneous action potentials: events were recorded with no clamp applied. For analysis, a positive-going threshold search was applied from baseline to 0 mV, and any events that crossed this upper threshold were included for further analysis. All AP measurements were made in Clampfit, then exported to excel where they were organized before being exported to Prism.

For I-step recordings: inclusion criteria included a steady baseline centered around the holding voltage, and action potential elicitation. For the rheobase, the average voltage of the step which first elicited an action potential was determined. For normalized rheobase, each rheobase voltage was divided by the capacitance of the cell in excel before being ultimately averaged together in Prism. Series resistance was measured in Clampfit, using steady sections of the sweeps from - 50 mV to 0 mV, then averaged in excel and exported to Prism.

For sodium current analysis: the trough of the first elicited event was measured against the baseline for each sweep, then averaged by voltage step and genotype before being exported to Prism.

For potassium currents: a flat section of the trace for all sweeps was measured for resistance using clampfit. These values were then exported to Excel, averaged by voltage step and genotype, and then exported to Prism.

For paired-pulse ratio analysis: the amplitude of the first response after each stimulus was measured, with each cell and recording having 7 replicate sweeps. These were averaged together by genotype and pulse width, then exported to Prism. Statistical analysis for PPR included two-tailed Student’s T-Tests for each pulse width, as well as two-way ANOVA with Tukey’s post hoc test. The amplitudes of every event from every first stimulation were pooled together by condition and averaged for the first response amplitudes.

For cumulative probability: first the inter-event-interval and absolute amplitude was calculated as described above. For the first events, the event start time was subtracted from the analysis window start time, then subsequent event start times were subtracted from the previous event start time in chronological order. These were then exported into a separate excel spreadsheet, where the values were binned (0.1 second bins for IEI, 2 pA bins for amplitude) and the cumulative frequency was calculated by summing the bins and dividing them by the total sum of all bins. These relative frequency values ranging from 0-20 seconds for IEI and from 0-200 -pA for amplitude were then exported to prism for graphing and analysis. A two-tailed Kolmogorov-Smirnov test was performed to determine significant differences in the distribution of these data.

### Morphology and synaptic puncta data collection

Coverslips were fixed 49 dpi and stained for VGLUT1 (Synaptic Systems, 135302), VGAT (Millipore, AB5062P), with TetO-EGFP expressed from the cells themselves. Cells were washed of media three times with DPBS (Gibco), fixed for 15 minutes with 4% Paraformaldehyde (Sigma), permeabilized with 0.1% Triton-X 100 (Sigma), and blocked with 5% Goat serum (Sigma) for 1 hour at room temperature and with gentle agitation. Primary antibodies were applied at a 1:500 dilution in blocking solution overnight at 4°C with gentle agitation. The coverslips were then washed three times with DPBS, and secondary antibodies (mouse 647: Thermo Fisher, rabbit 546: Thermo Fisher) were applied at a 1:1000 dilution in blocking solution for one hour at room temperature with gentle agitation. After the final wash, coverslips were mounted to glass slides with Fluoromount (Southern Biotech) and imaged at 20x magnification for neurite morphology and 60x magnification (with 2x digital zoom) for synaptic puncta on a Nikon A1R confocal microscope (Nikon). Morphology was analyzed by tracing the GFP-positive cells in Imaris 10.1 (Oxford Instruments). Puncta analysis was performed using the 60x images in Imaris. A 10 µm segment of dendrite was isolated and analyzed for colocalization with VGLUT1 or VGAT, excluding any signal more that 1 µm from the dendrite surface.

### Image quantification of neurite outgrowth and synaptic puncta analysis

Images taken as described in the “morphology and puncta acquisition” section were blinded to the analyzer and randomly ordered, then imported to Imaris image analysis software. Here, images were analyzed as described here: for morphological analysis, LUTs were adjusted such that GFP was clearly identifiable. Cells with too dim or blurred of an outline were rejected. Filamenting and surface creation tools in Imaris were used to outline the cell and dendrites, and export relevant metrics for analysis.

For puncta analysis, the 60x images were imported to Imaris, and the filamenting tools were used to measure distance along a measured dendrite. In parallel, surfaces were created for each channel (GFP (dendritic section) VGAT and VGLUT (whole image)) with the following settings: Surface detail: 0.3 µm; Background subtraction: 1.00 µm; Seed point diameter: 0.3 µm. Also, a filter for the number of voxels (>10), and surface objects farther than 0.1 µm from the dendrite were filtered out. Dendrites as close to 150 µm were chosen for analysis, but actual dendritic segment lengths varied greatly. The number of puncta for each channel along the analyzed dendrite segment were counted, then divided by the total dendritic segment length and multiplied by ten to calculate the average puncta density (reported as puncta/10 µm). Puncta volume was also determined by carefully adjusting the gating and seed point density for breaking up conjoined surface objects (varied by image) such that doublets or triplets were reduced, then exporting the average volume values to excel for tidying, then ultimately Prism for graphing and analysis. Statistical analysis was performed using a Student’s T-Test; nonsignificant comparisons not included.

## Data and code availability

This paper does not report original code.

## Acknowledgements

This work was supported by NIMH (R01MH122519 to C.P.), UMass Amherst IALS (Translational fellowship to J.E.), NIGMS BTP training grant (T32GM135096 to J.E.). We thank SFARI for generation of iPSCs from ASD patients. We thank members of the Pak and Pang labs for support and discussion on the project.

## Author information

C.P., Z.P., and J.E. designed the study, analyzed the data, and wrote the manuscript. T.S. and S.J.G. designed and executed functional connectivity analysis. J.E. and D.M. performed whole cell patch clamp electrophysiology and analysis. J.E. and E.H. performed HD-MEA recordings. J.E. performed experiments and analysis of morphology and synaptic puncta. F.R., R.S., and D.M. assisted with lentivirus generation and cell culture. M.H., L.P., and L.W. assisted with experiments.

