## Supplementary Figures and Text for "Distinct Synaptic Mechanisms Drive *NRXN1* Variant-Mediated Pathogenesis in iPSC-Derived Neuronal Models of Autism and Schizophrenia"

**Figure S1 sPSC analysis individual pairs**

**Figure S2 neurite outgrowth analysis individual pairs**

**Figure S3 synapse density analysis individual pairs**

**Figure S4 mEPSC analysis individual pairs**

**Figure S5 mIPSC analysis individual pairs**

**Figure S6 MEA control experiment**

**Figure S7 functional connectivity analysis in individual pairs**

**Figure S8 mini analysis in second line of SCZ-NRXN1 deletion**

**Figure S9 mini analysis in NRXN1 cKO**

**Supp Table 1 iPSC line description**

**Supp Table 2 Statistical analysis summary**

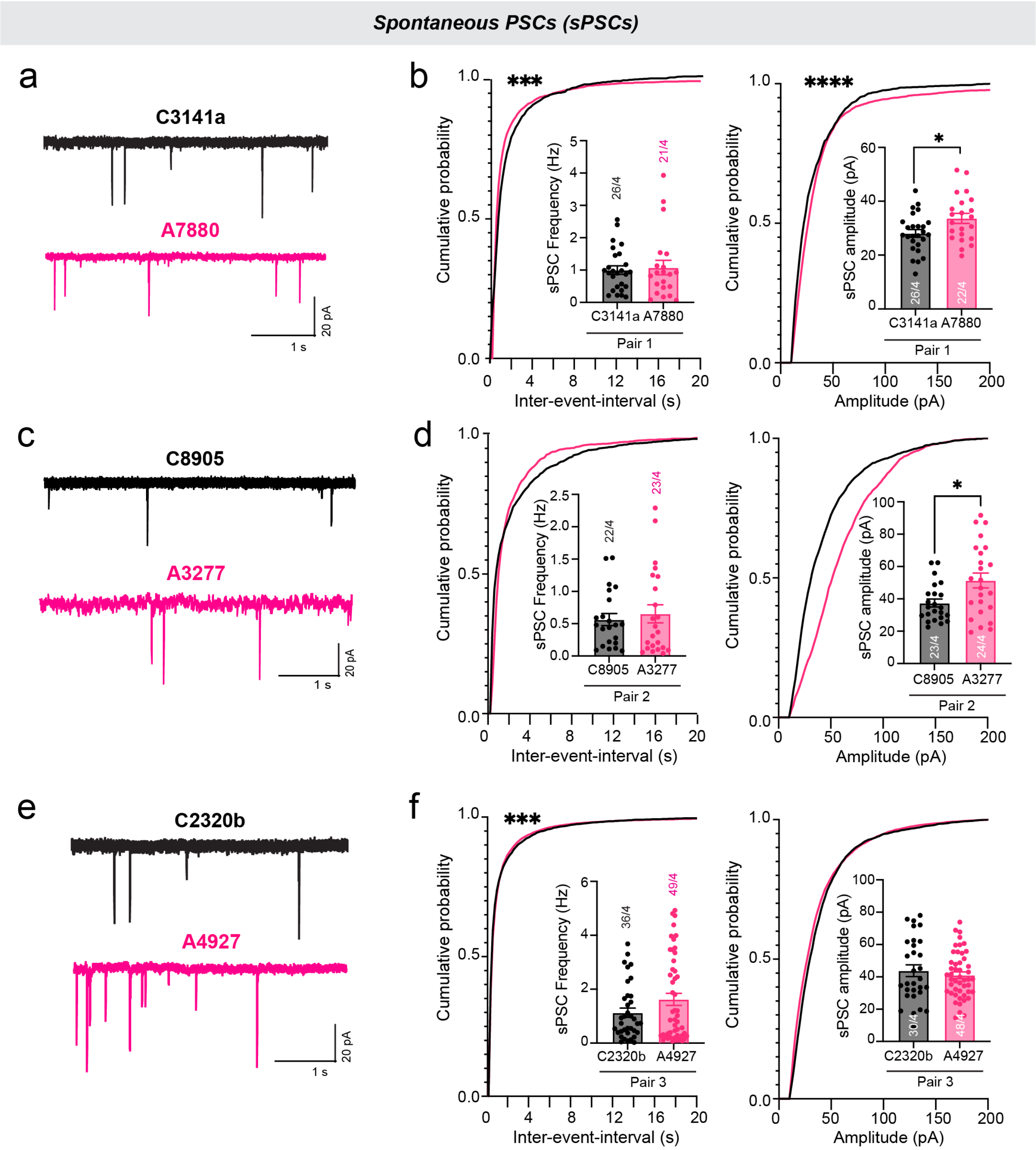


**Figure S1 (related to Figure 3). Spontaneous postsynaptic current analysis of individual pairs. a, c, e)** Representative sPSC traces (left column) and quantifications of sPSC frequency and amplitude (right column) of each pair – **b)** pair 1, **d)** pair 2, and **f)** pair 3.

All summary data are means +/- SEM (numbers inside or above bars show number of cells/independent cultures analyzed). Cells were recorded at 42-49 dpi. Statistical significance for average frequency and amplitude were determined with Student's T-Test, whereas significance for cumulative probability distributions was determined by two-tailed Kolmogorov-Smirnov test. P-values: p* < 0.05, p***<0.001, p****<0.0001.

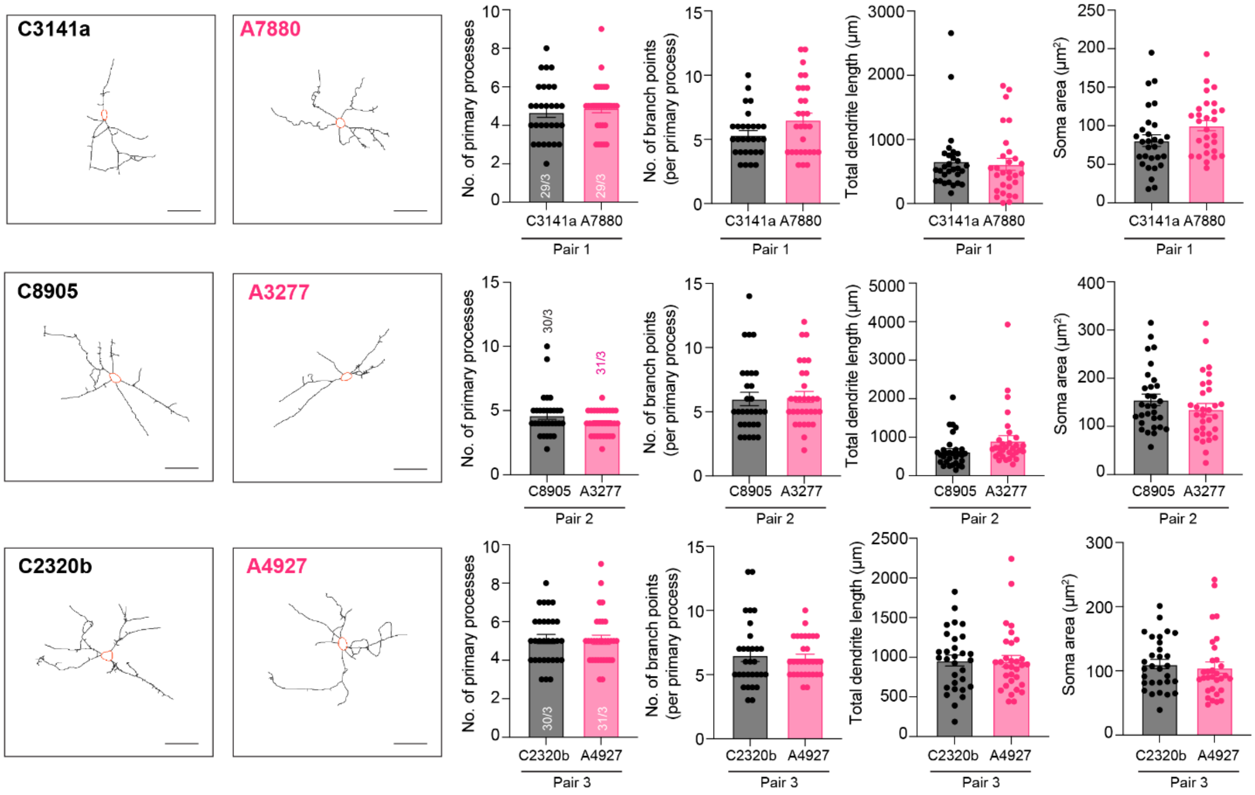


**Figure S2 (related to Figure 4). Individual pair analysis of neuronal morphology.**

**Left:** Representative binary mask of EGFP+ excitatory iNs in the E-I mixed cultures (scale bar, 40 um). **Right:** Quantification summary graphs of various parameters of neuronal morphology (number of primary processes, number of branch points per primary processes, total dendritic length, and soma size) in control vs. ASD iNs.

All summary data are means +/- SEM (numbers in bars or parentheses show number of cells/independent cultures analyzed). Cells were fixed for analysis on dpi 49. No significant differences were detected in any of the conditions by Student’s t-test. Nonsignificant comparisons are not indicated.

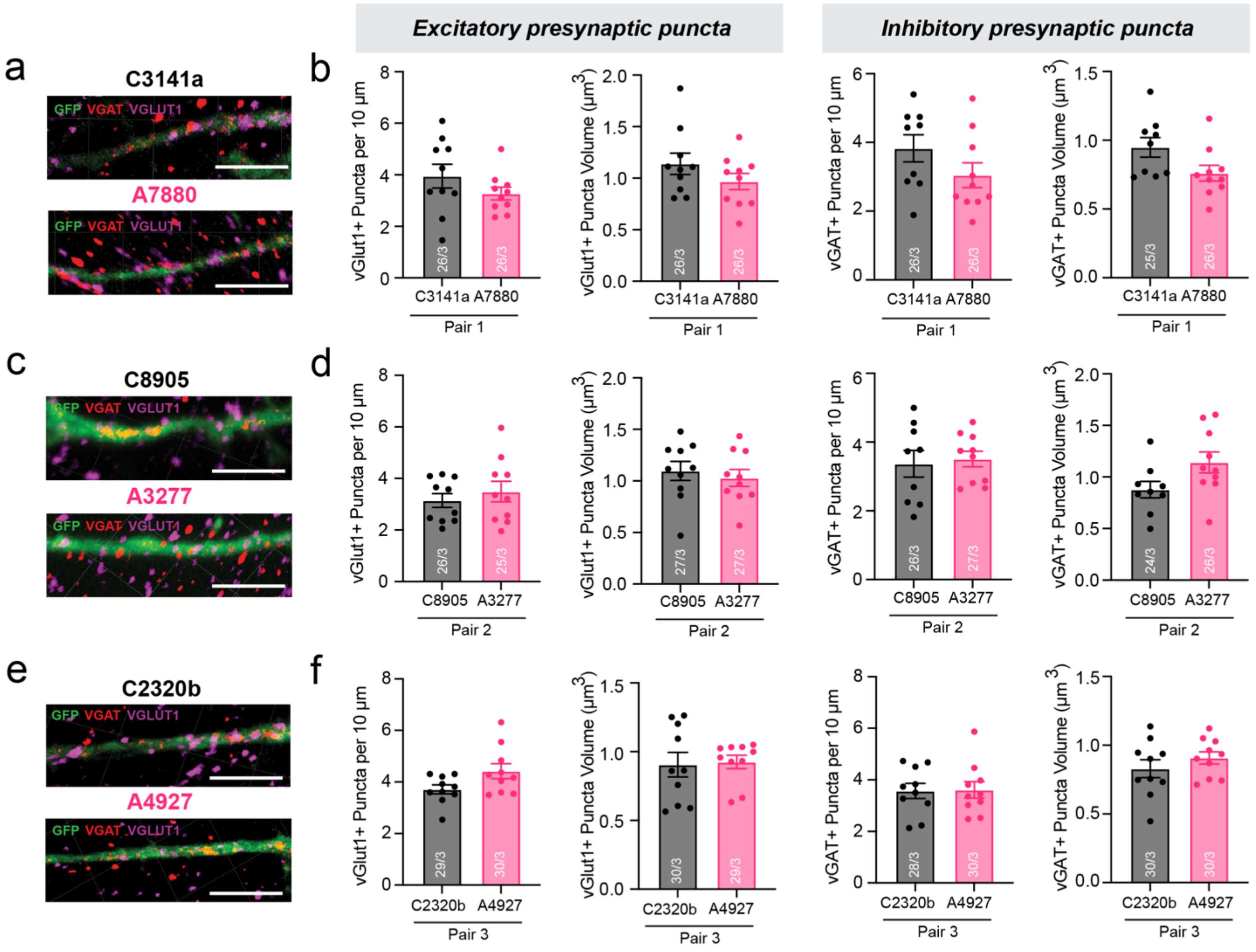


**Figure S3 (related to Figure 4). Individual pair analysis of synaptic puncta density and size.**

**a, c, e)** Representative images of isolated EGFP+ excitatory neuronal dendritic segments stained for vGlut1 (excitatory synapse marker) and vGAT (inhibitory synapse marker) (scale bar, 5 um). **b ,d, f)** Quantification summary graphs of the puncta density and volume in control vs. ASD iNs.

All summary data are means +/- SEM (numbers in bars or parentheses show number of cells/independent cultures analyzed). Cells were fixed for analysis on dpi 49. No significant differences were detected in any of the conditions by Student's t-test. Nonsignificant comparisons are not indicated.

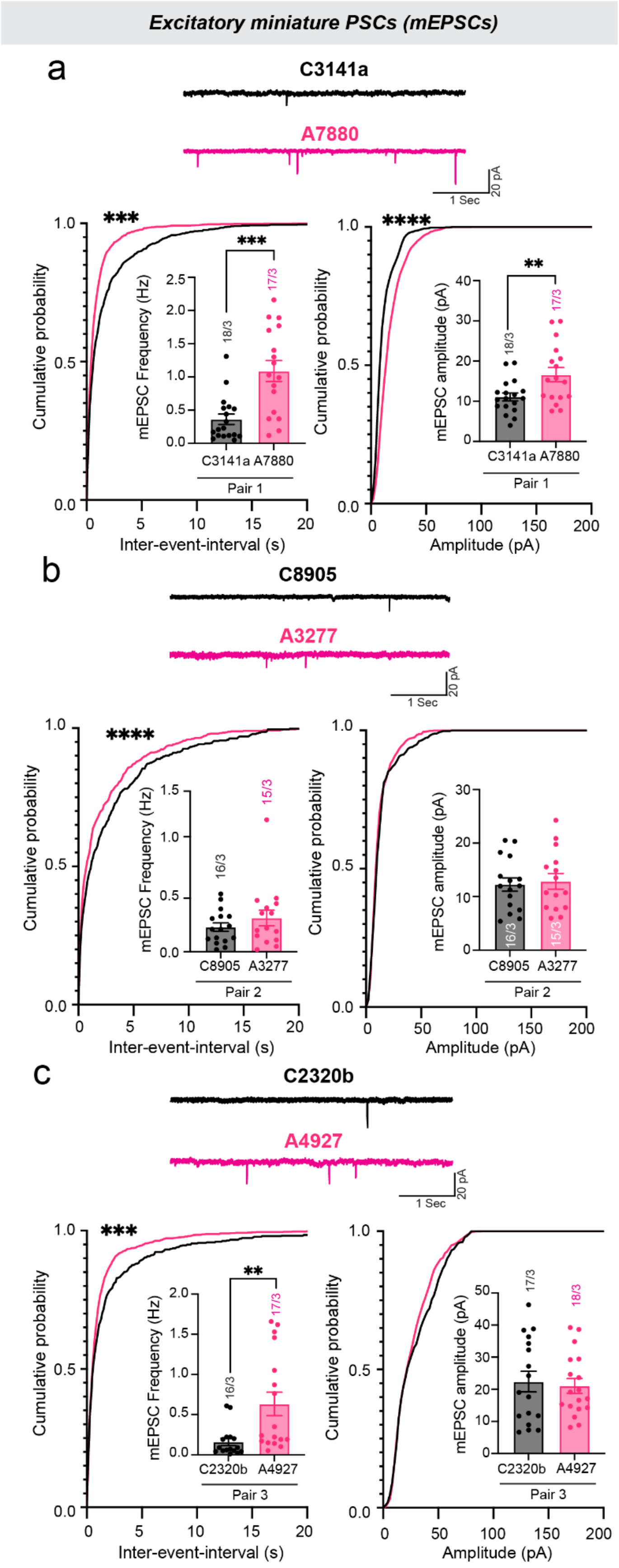


**Figure S4 (related to Figure 4). Analysis of spontaneous excitatory miniature synaptic transmission in pairs.** Representative mEPSC traces (top) and quantifications of mEPSC frequency and amplitude (bottom) of each pair – **a)** pair 1, **b)** pair 2, and **d)** pair 3. Recordings were performed in the presence of TTX (1 uM) to block all APs to capture single vesicular release miniature events and in PTX (50 uM) to inhibit GABAergic inhibitory postsynaptic currents. Quantifications are shown both as cumulative probability plots and as bar graphs (inserts).

All summary data are means +/- SEM (numbers in bars or parentheses show number of cells/independent cultures analyzed). Cells were recorded at 42-49 dpi. Statistical significance for average frequency and amplitude were determined with Student's T-Test, whereas significance for cumulative probability distributions was determined by two-tailed Kolmogorov-Smirnov test. P-values: ** < 0.01, ***<0.001, ****<0.0001.

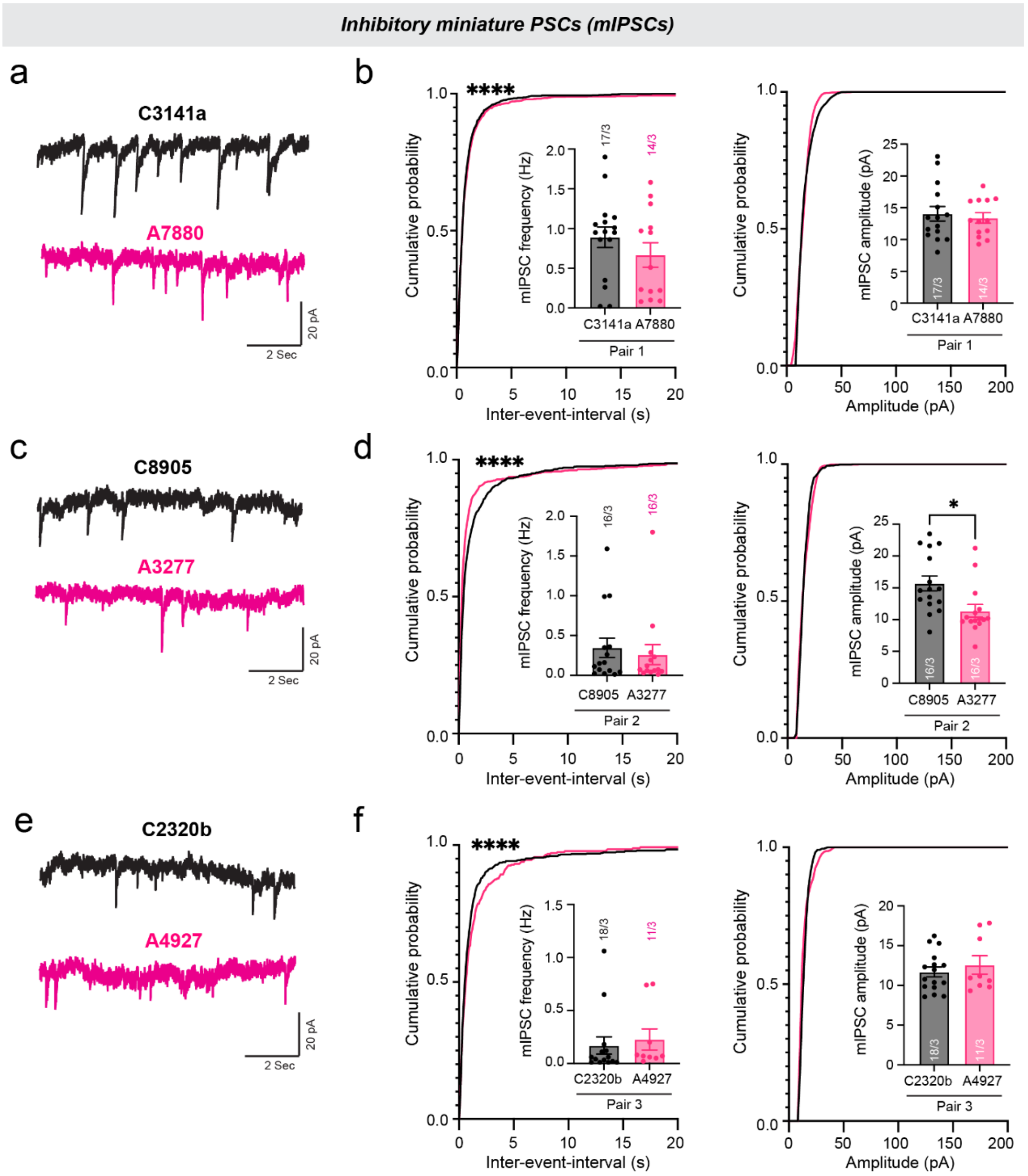


**Figure S5 (related to Figure 4). Analysis of spontaneous inhibitory miniature synaptic transmission in pairs. a, c, e)** Representative mIPSC traces (left column) and quantifications of mIPSC frequency and amplitude (right column) of each pair – **b)** pair 1, **d)** pair 2, and **f)** pair 3. Recordings were performed in the presence of TTX (1 uM) to block all APs to capture single vesicular release miniature events and in CNQX (20 μM)/APV (50 μM) to inhibit AMPAR- and NMDAR-mediated glutamatergic postsynaptic currents. Quantifications are shown both as cumulative probability plots and as bar graphs (inserts).

All summary data are means +/- SEM (numbers in bars or parentheses show number of cells/independent cultures analyzed). Cells were recorded at 49-56 dpi. Statistical significance for average frequency and amplitude were determined with Student's T-Test, whereas significance for cumulative probability distributions was determined by two-tailed Kolmogorov-Smirnov test. P-values: ** < 0.01, ****<0.0001.

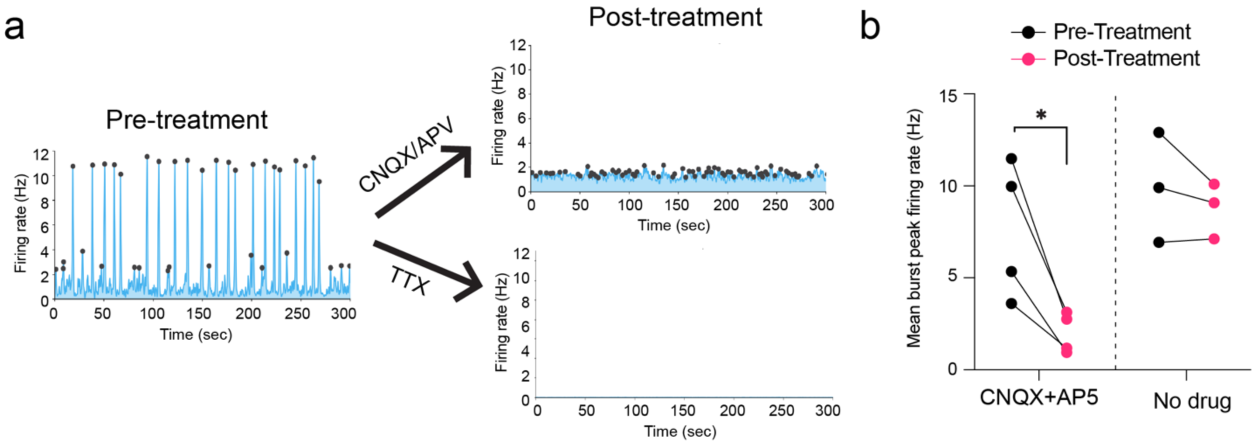


**Figure S6 (related to Figure 6). Neuronal spikes and network bursts are synaptically driven responses. a)** Firing rate graphs showing the effects of glutamate receptor blockers (CNQX, 20 µm; APV, 50µm) and sodium channel blocker (TTX, 1 µm) on neuronal networks pre- and post-treatments. **b)** Summary graphs of grouped average burst peak firing rate (Hz) pre- and post-treatments (repeated measures). No drug condition was used as a negative control. N=4 for CNQX/APV and N=3 for No drug.

Control iPSC-Ngn2-iNs were differentiated and plated on HD-MEA chips at 3 dpi and cultured for an additional 32 days. Prior to treatment with blockers, 5-min recordings of neuronal spike activity and network bursting were performed and analyzed (pre-treatment). Immediately following treatment (post-treatment), a 30-min network recording was performed to extract neuronal spike and burst level data. Paired Student's t-test; *$p<0.05$.

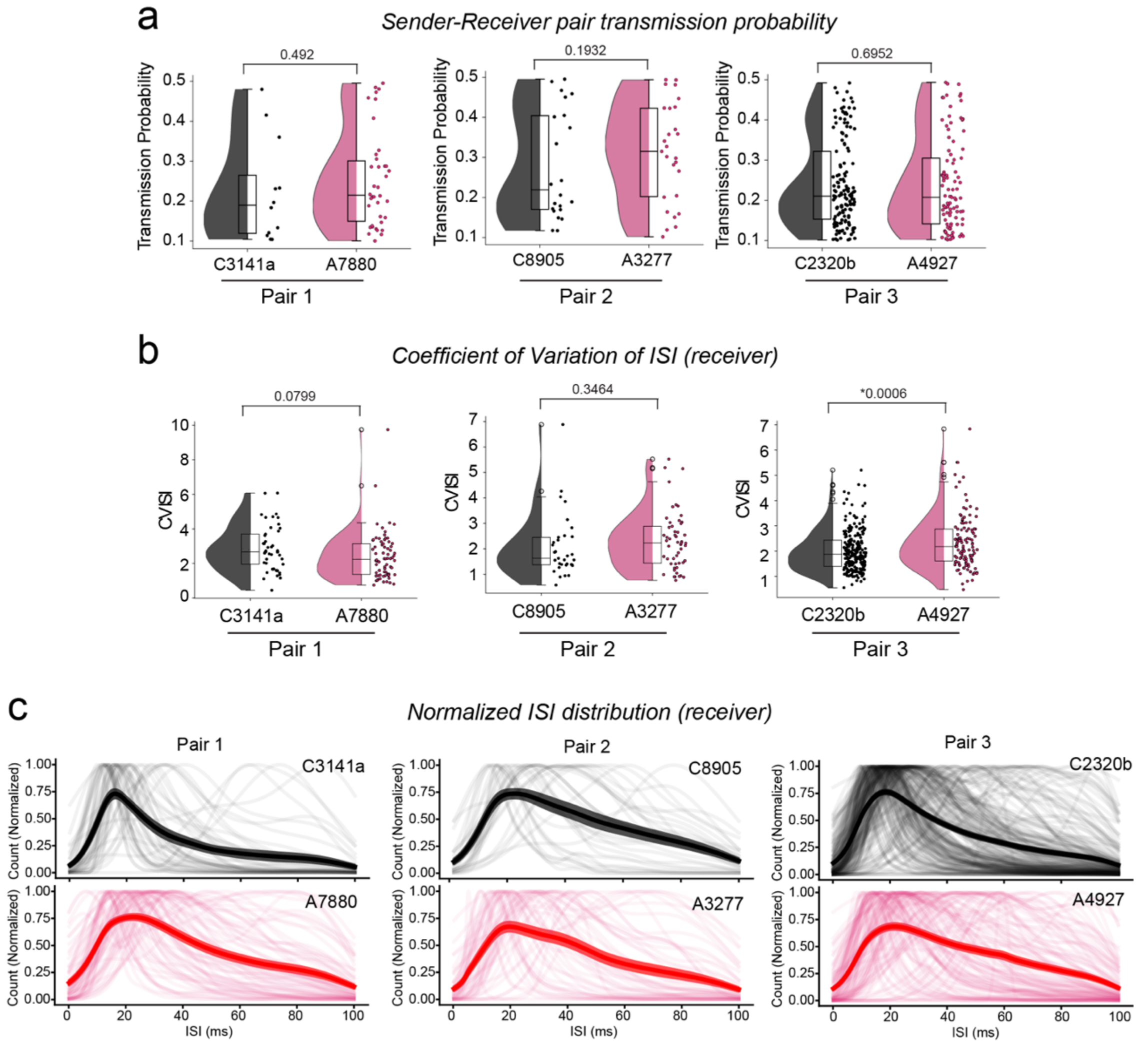


**Figure S7 (related to Figure 6) Functional connectivity analysis from NRXN1+/- ASD patient iPSCs.** HD-MEA recordings obtained from 28 dpi Ngn2 solo cultures from 3 $\mathit{NRXN1}^{+/-}$ ASD vs. 3 control iPSC lines (n>6 independent cultures per line). Units displaying functional connectivity across distances on the HD-MEA. Units are spike sorted, and functional connectivity is determined using spike train cross-correlation. **a)** Pre- and post-synaptic spike transmission probability in *Ngn2* cultures from each pair of three control and three ASD genotypes. Box plots showing median values of transmission probability with upper quartile (75%) and lower quartile (25%). Increased transmission probability indicates increased likelihood that sender and receiver are functionally connected. **b)** Coefficient of variation (CV) of interspike intervals (ISI) for receivers in *Ngn2* cultures, with pair analysis of the three control and three ASD genotypes. Box plots showing median values of CV ISI with upper quartile (75%) and lower quartile (25%). Increased CV ISI indicates more irregular firing patterns. **c)** Normalized ISI distribution for receivers in each paired Ngn2 culture. Distribution is plotted for ISI in the range of [0, 100] ms, smoothened using kernel density estimation (KDE) with a Gaussian kernel and normalized by maximum value. The thick line with a shaded region on the graph represents the mean and SEM.

Statistical significance for transmission probability and CV ISI was determined with Student's T-Test and p-values are indicated on the top of each plot.

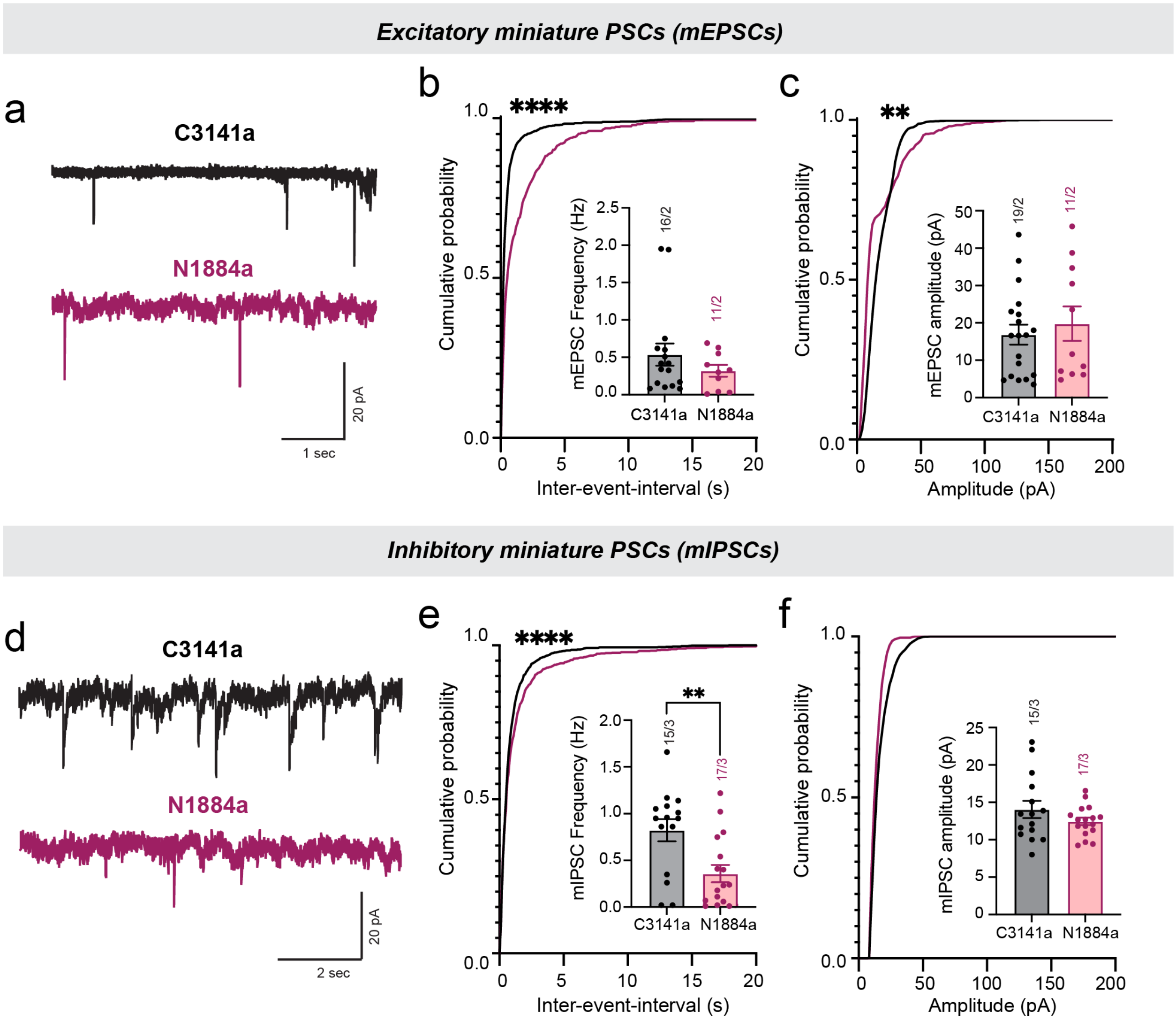


**Figure S8 (related to Figure 7). A second line of *NRXN1*$^{+/-}$ SCZ patient-derived E-I iNs show a similar decrease in synaptic transmission.** Representative mEPSC **(a)** and mIPSC traces **(d)** and quantifications of their frequency and amplitude (mEPSC – **b, c**; mIPSC – **e, f**) in N1884a, an independent iPSC line derived from SCZ patient carrying NRXN1 deletion, and the same C3141a control iPSC line used for Figure 7.

All summary data are means +/- SEM (numbers in bars or parentheses show number of cells/independent cultures analyzed). Cells were recorded at 42-49 dpi for mEPSCs and 49-56 dpi for mIPSCs. Statistical significance for average frequency and amplitude were determined with Student's T-Test, whereas significance for cumulative probability distributions was determined by two-tailed Kolmogorov-Smirnov test. P-values: *p<0.01, ****p<0.0001.

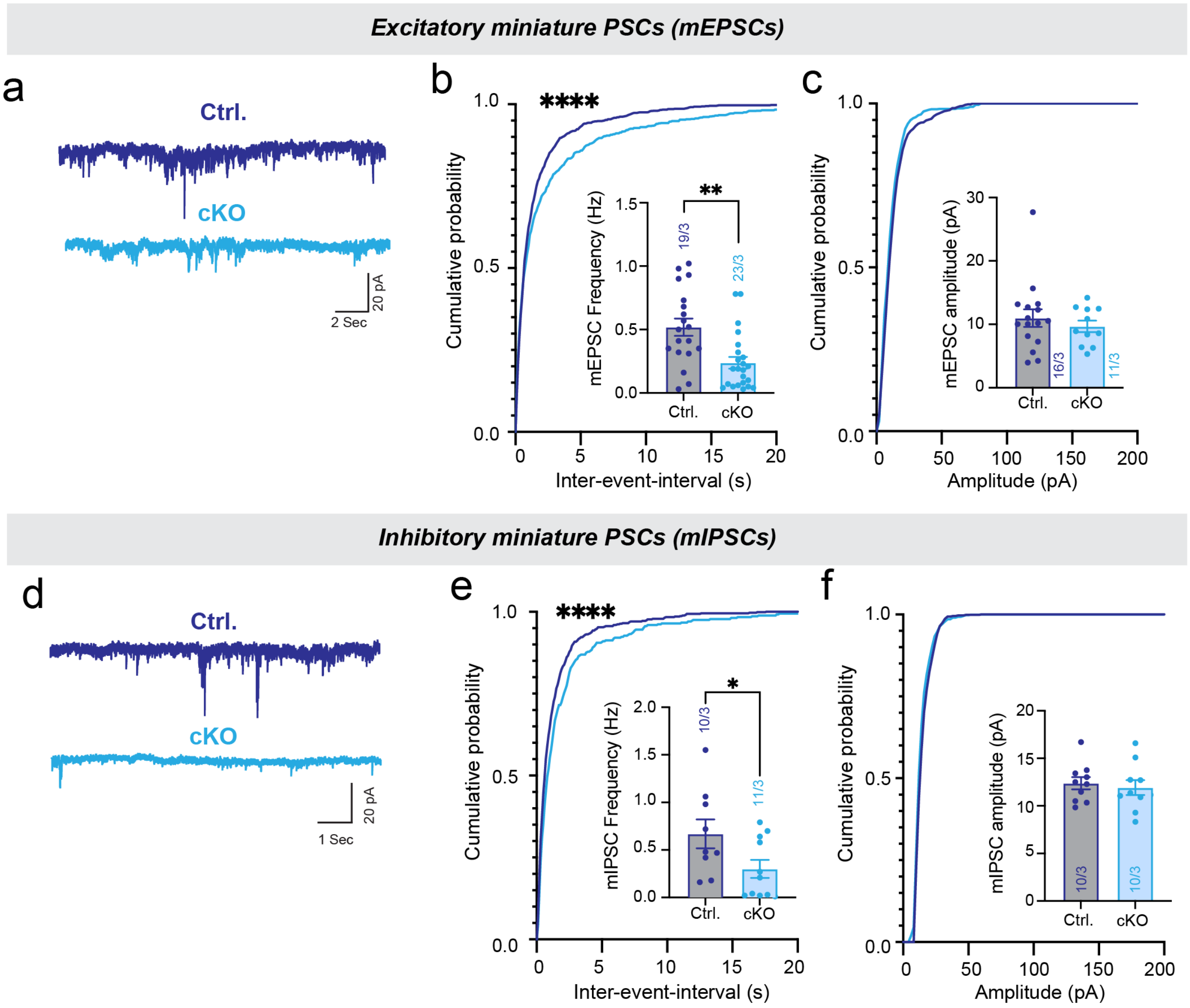


**Figure S9 (related to Figure 7). NRXN1 heterozygous cKO E-I iNs also show decreased excitatory and inhibitory miniature frequencies similar to *$NRXN1^{+/-}$* SCZ patient-derived E-I iNs. a)** Representative mEPSC traces and quantifications of mEPSC frequency **(b)** and amplitude **(c)** in Ctrl (Flp) and cKO (Cre)**. d)** Representative mIPSC traces and quantifications of mIPSC frequency **(e)** and amplitude **(f)** in Ctrl (Flp) and cKO (Cre)**.**

All summary data are means +/- SEM (numbers in bars or parentheses show number of cells/independent cultures analyzed). Cells were recorded at 42-49 dpi for mEPSCs and 49-56 dpi for mIPSCs. Statistical significance for average frequency and amplitude were determined with Student's T-Test, whereas significance for cumulative probability distributions was determined by two-tailed Kolmogorov-Smirnov test. P-values: *p< 0.05, **p<0.01,****p<0.0001.

**Supplementary Table 1. Genomic coordinates of *NRXN1+/-* deletion breakpoints in iPSCs**

| Line ID | Sex | NRXN1 deletion | Deletion start-stop (hg38) | Deletion size (bp) | Diagnosis | Publication |
|---|---|---|---|---|---|---|
| C3141a | M | Control | - | - | None | Pak et al., 2021 |
| C8905a | M | Control | - | - | None | Pak et al., 2021 |
| C2320b | M | Control | - | - | None | Pak et al., 2021 |
| A7880 | M | Exon 1-2 | Chr2: 51,020,156-51,084,394 | 64,239 | ASD | Unpublished; this study |
| A4927 | M | Exon 14 PTV | Chr 2: 50,497,467 | A-to-T | ASD | Unpublished; this study |
| A3277 | M | Exon 1-2 | Chr 2: 51,018,339-51,106,818 | 88,480 | ASD | Unpublished; this study |
| N3320a | M | Exon 1-6 | Chr2:50,899,162-51,081,962 | 182,801 | SCZ | Pak et al., 2021 |
| N1884a | M | Exon 1-2 | Chr2:50,979,562-51,087,962 | 108,400 | SCZ | Pak et al., 2021 |
